# Cell Type- and Tissue-specific Enhancers in Craniofacial Development

**DOI:** 10.1101/2023.06.26.546603

**Authors:** Sudha Sunil Rajderkar, Kitt Paraiso, Maria Luisa Amaral, Michael Kosicki, Laura E. Cook, Fabrice Darbellay, Cailyn H. Spurrell, Marco Osterwalder, Yiwen Zhu, Han Wu, Sarah Yasmeen Afzal, Matthew J. Blow, Guy Kelman, Iros Barozzi, Yoko Fukuda-Yuzawa, Jennifer A. Akiyama, Veena Afzal, Stella Tran, Ingrid Plajzer-Frick, Catherine S. Novak, Momoe Kato, Riana D. Hunter, Kianna von Maydell, Allen Wang, Lin Lin, Sebastian Preissl, Steven Lisgo, Bing Ren, Diane E. Dickel, Len A. Pennacchio, Axel Visel

**Author notes:** To whom correspondence should be addressed: A.V. Current address.

## Abstract

The genetic basis of craniofacial birth defects and general variation in human facial shape remains poorly understood. Distant-acting transcriptional enhancers are a major category of non-coding genome function and have been shown to control the fine-tuned spatiotemporal expression of genes during critical stages of craniofacial development^1–3^. However, a lack of accurate maps of the genomic location and cell type-specific *in vivo* activities of all craniofacial enhancers prevents their systematic exploration in human genetics studies. Here, we combined histone modification and chromatin accessibility profiling from different stages of human craniofacial development with single-cell analyses of the developing mouse face to create a comprehensive catalogue of the regulatory landscape of facial development at tissue- and single cell-resolution. In total, we identified approximately 14,000 enhancers across seven developmental stages from weeks 4 through 8 of human embryonic face development. We used transgenic mouse reporter assays to determine the *in vivo* activity patterns of human face enhancers predicted from these data. Across 16 *in vivo* validated human enhancers, we observed a rich diversity of craniofacial subregions in which these enhancers are active *in vivo*. To annotate the cell type specificities of human-mouse conserved enhancers, we performed single-cell RNA-seq and single-nucleus ATAC-seq of mouse craniofacial tissues from embryonic days e11.5 to e15.5. By integrating these data across species, we find that the majority (56%) of human craniofacial enhancers are functionally conserved in mice, providing cell type- and embryonic stage-resolved predictions of their *in vivo* activity profiles. Using retrospective analysis of known craniofacial enhancers in combination with single cell-resolved transgenic reporter assays, we demonstrate the utility of these data for predicting the *in vivo* cell type specificity of enhancers. Taken together, our data provide an expansive resource for genetic and developmental studies of human craniofacial development.

**Graphical Abstract:** 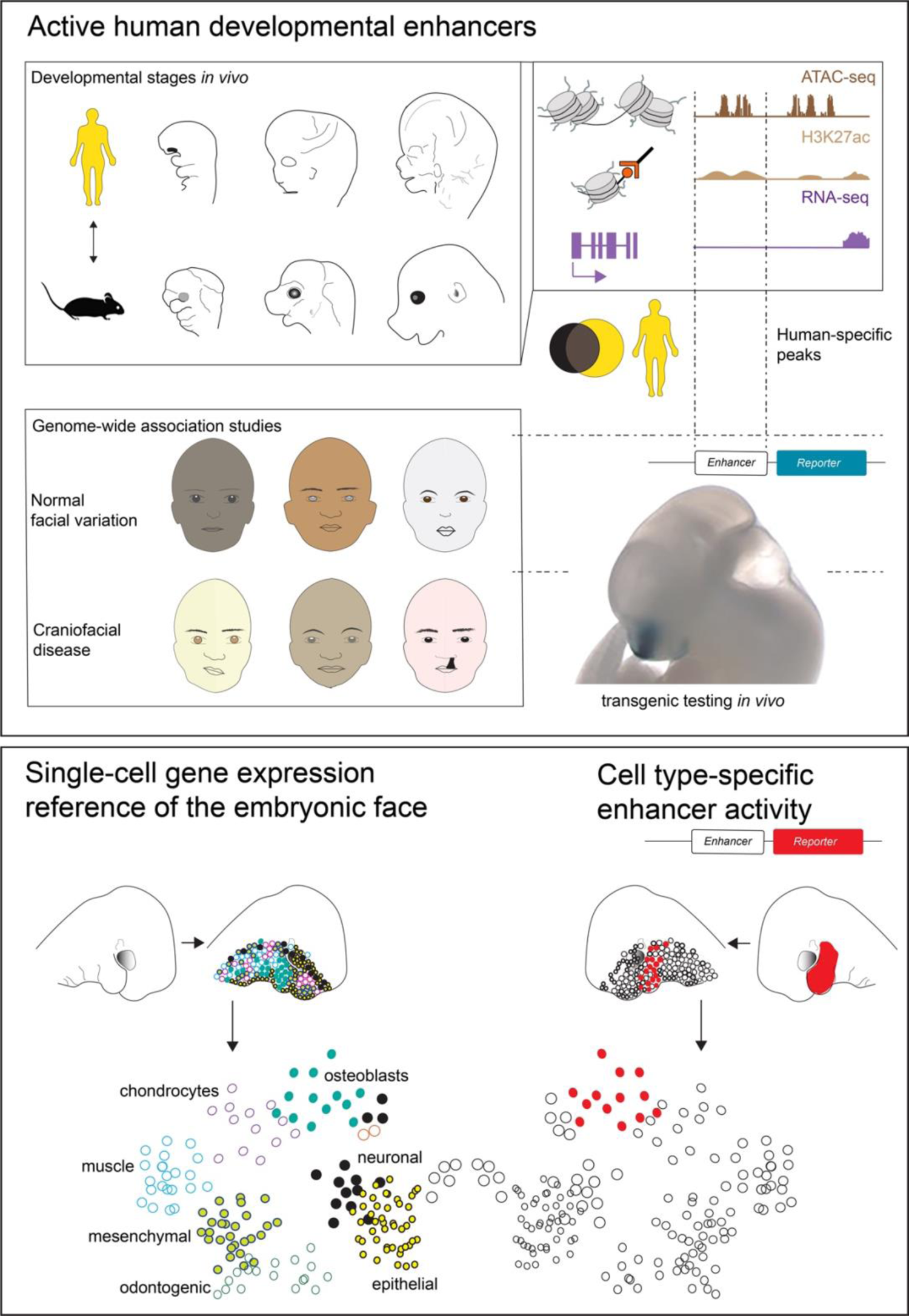

## Introduction

The development of the human face is a highly complex morphogenetic process. It requires the precise formation of dozens of intricate structures to enable the full complement of facial functions including food uptake, breathing, speech, major sensory functions including hearing, sight, smell, taste, and nonverbal communication through facial expression. Intriguingly, these functional constraints coincide with substantial inter-individual variation in facial morphology, which humans use as the principal means for recognizing each other. Apart from providing the basis for normal facial variation, early developmental processes underlying facial morphogenesis are highly sensitive to genetic abnormalities as well as environmental effects^4^. Even subtle disturbances during embryogenesis can result in a range of craniofacial defects or dysfunctions^5^.

In embryonic facial development, the primary germ layers as well as the neural crest contribute crucially to the formation of the pharyngeal arches, the frontonasal process and the midface, which in combination give rise to the derived structures of the face^6–9^. The primary palate forms by the fifth week post conception^10^ and development of primary palate derivatives, secondary palate, and many other structures, combined with overall rapid growth, result in a discernable human-like appearance by the tenth week post conception^11^. Genetic or environmental perturbations during these crucial developmental stages are known to result in craniofacial malformations of varying severity and of typically irreversible nature^12–16^. Development of the mammalian face requires a conserved set of genes and signaling pathways^17^, which are regulated by distant-acting transcriptional enhancers that control gene expression in time and space^1, 18–24^. Together with the genes they control, these enhancers are a critical component of mammalian craniofacial morphogenesis. It is estimated that there are hundreds of thousands of enhancers in the human genome for approximately 20,000 genes^25^ and chromatin profiling studies have identified initial sets of enhancers predicted to be active in craniofacial development^1, 25, 26^. However, these data sets do not cover critical stages of human facial development, such as secondary palate formation, and provide no information about the cell type specificity of individual enhancers. In part due to the continued incomplete annotation state of the craniofacial enhancer landscape, the number of enhancers that could be mechanistically linked to facial variation or craniofacial birth defects has remained limited^1, 18–23^. With an increasingly refined view of the genetic variation underlying human facial variation^27^ and whole genome sequencing as an increasingly common clinical approach for the identification of noncoding mutations in craniofacial birth defect patients^28, 29^, an expanded and accurate map of human craniofacial enhancers is critical for interpretation of any noncoding findings emerging from these studies. Here we provide a comprehensive compilation of regulatory regions from the developing human face during embryonic stages critical for birth defects including orofacial clefts, along with cell type-specific gene expression and open chromatin signatures for the developing mammalian face.

## Results

### Epigenomic Landscape of the Human Embryonic Face

To map the epigenomic landscape of critical periods of human face development, we focused on Carnegie stages (CS) 18-23, a period coinciding with the formation of important structures including the maxillary palate, rapid overall growth, and significant changes in the relative proportions of craniofacial structures that impact on ultimate craniofacial shape^11, 30, 31^. These stages are of direct clinical relevance because common craniofacial defects, including cleft palate and major facial dysmorphologies, result from disruptions within this developmental window (**Figure 1a**)^32, 33^. To determine the genomic location of enhancers, we generated genome-wide maps of the enhancer-associated histone mark H3K27ac (ChIP-seq), accessible chromatin (ATAC-seq), and gene expression (RNA-seq) from embryonic face tissue for CS18, 19, 22, and 23 (**Supplemental Table 1**). To extend our compendium to earlier stages, we complemented this data with published H3K27ac peaks (ChIP-seq) from CS13-17 human face tissue^26^ (**Supplemental Table 1, Methods**). In total, we observed 13,983 reproducible human candidate enhancers, as defined by the presence of H3K27ac signal in at least two biological samples at any stage between CS13-23 of development (**Supplemental Table 2**). Of the 10,893 regions marked by H3K27ac at stages CS18-23, 6,718 (61.7%) also showed accessible chromatin signal, further supporting their expected status as active enhancers (**Supplemental Table 3, Methods)**.

**Figure 1.**
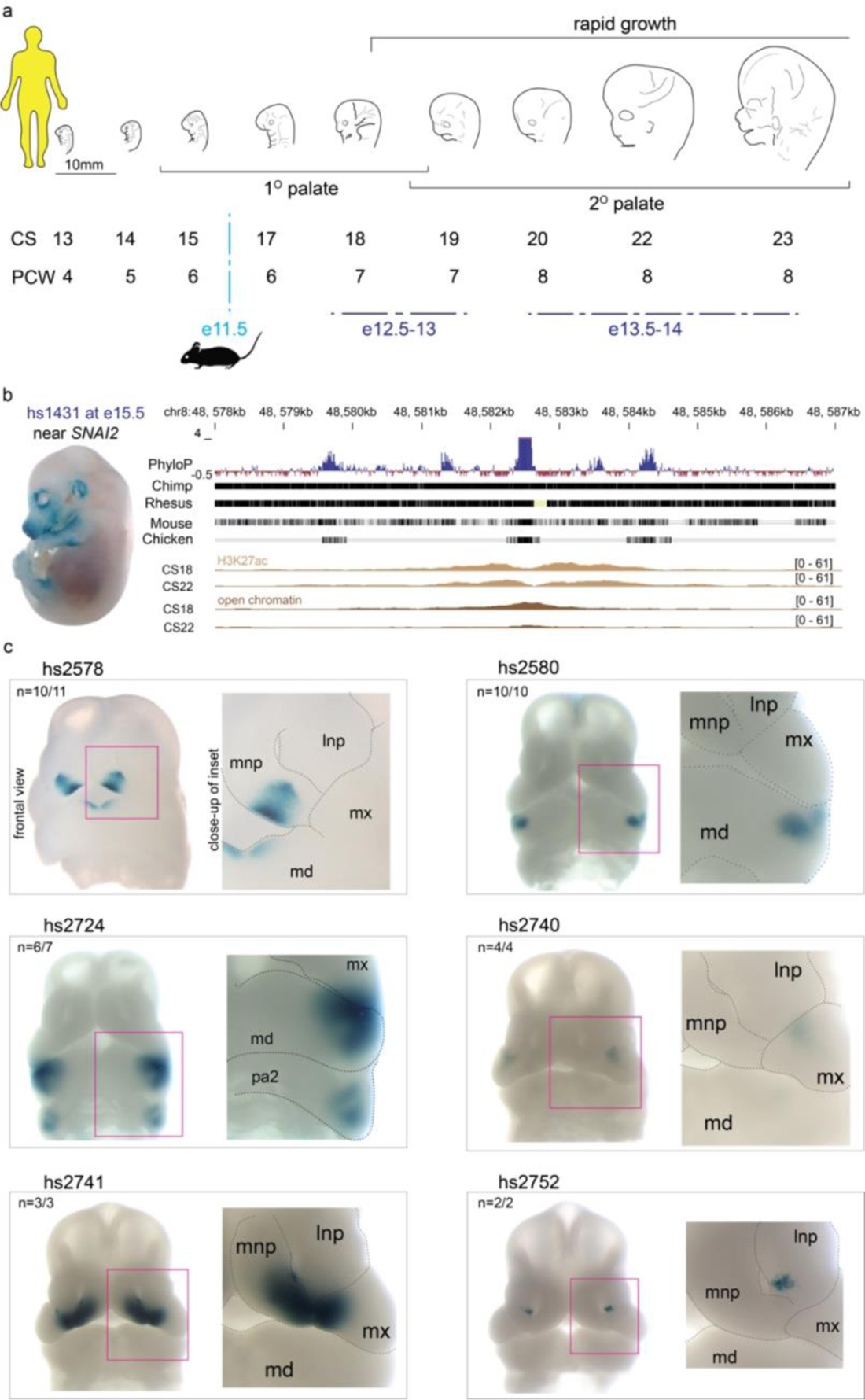
Developmental enhancers in human craniofacial morphogenesis. **a.** Developmental time points coinciding with critical windows of craniofacial morphogenesis are shown by Carnegie stage (CS) and post-conceptional week (PCW) in humans, and comparable embryonic (e) stages for mouse are shown in embryonic days. **b.** Representative embryo image at e15.5 for an *in vivo* validated enhancer (hs1431) shows positive *lacZ*-reporter activity in craniofacial structures (and limbs). Adjacent graphic shows the genomic context and evolutionary conservation of the region, with H3K27ac-bound and open chromatin regions located within the hs1431 element. **c.** Six examples of human craniofacial enhancers discovered in this study with *in vivo* activity validated in e11.5 transgenic mouse embryos. Enhancers hs2578, hs2580, hs2724, hs2740, hs2741 and hs2752 show *lacZ*-reporter activity in distinct subregions of the developing mouse face. Lateral nasal process (lnp), medial nasal process (mnp), maxillary process (mx), and mandibular process (md). n, reproducibility of each pattern across embryos resulting from independent transgenic integration events.

For an initial assessment of the biological relevance of this genome-wide set of predicted human craniofacial enhancers, we compared it with the large collection of *in vivo*-validated enhancers available through the VISTA enhancer browser^34^. Among the 3,193 elements that have been tested in VISTA to date, we identified 153 cases in which an enhancer predicted through the present human-derived epigenomic dataset had been previously shown to drive reproducible expression in embryonic facial structures including the nose, branchial arches, facial mesenchyme, cranial nerves, melanocytes, ear, or eye (**Extended Data Figure 1, Supplemental Table 4**). A representative example of a previously validated VISTA craniofacial enhancer is shown in **Figure 1b**.

To assess the value of these data for the discovery of additional craniofacial *in vivo* enhancers in the human genome, we tested 60 candidate human enhancers in a transgenic mouse assay (**Methods, Supplemental Table 5**). We identified 16 cases of previously unknown enhancers that showed reproducible activity in craniofacial structures. **Figure 1c** illustrates the rich diversity of craniofacial structures in which these enhancers drive reproducible *in vivo* activity. Examples include enhancers driving expression in restricted subregions of the medial nasal process and mandible (hs2578), the mandible (hs2580), the mandible and second pharyngeal arch (hs2724), the maxillary (hs2740), the medial nasal process and maxillary (hs2741), or the lateral nasal process (hs2752, **Fig. 1c**).

### Functional Associations, Developmental Dynamics, and Conservation of Human Craniofacial Enhancers

To further assess the biological relevance of the human candidate enhancer sequences identified by our approach, we examined known functions of their presumptive target genes using rGREAT ontology analysis^35^. The identified candidate enhancers are enriched near genes implicated in craniofacial human phenotypes, with 9 of the top 15 terms directly related to craniofacial or eye-associated phenotypes (**Figure 2a**), including midface retrusion, reduced number of teeth, and abnormality of maxilla.

**Figure 2.**
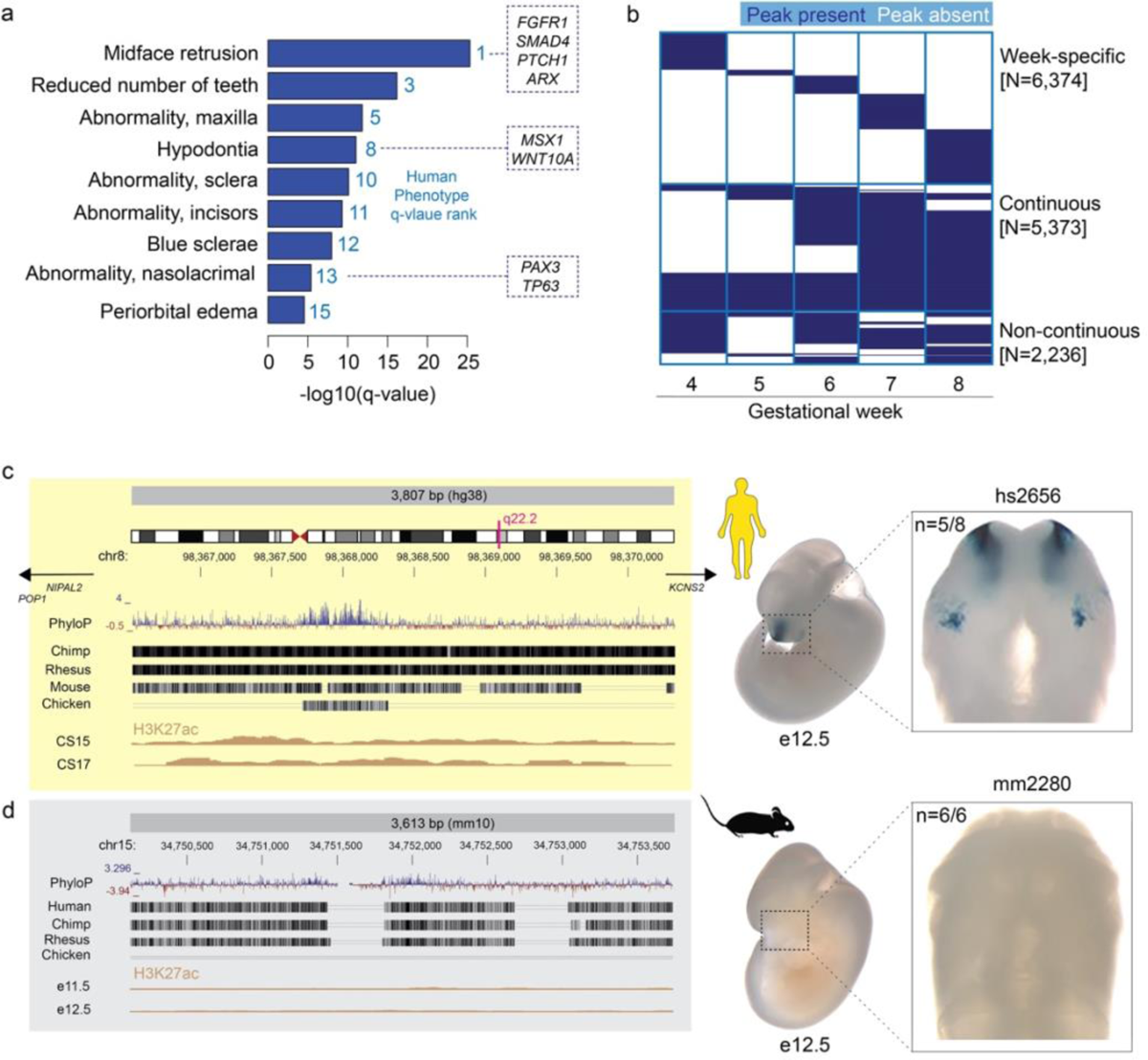
Developmental dynamics and conservation of human craniofacial enhancers. **a.** Results of rGREAT ontology analysis for 13,983 highly reproducible human craniofacial enhancers, ranked by Human Phenotype q-value. The ontology terms indicate that our predictions of human craniofacial enhancers are enriched near presumptive target genes known to play important roles in craniofacial development (examples in boxes). **b.** Predicted activity windows of 13,983 candidate human enhancers (rows) arranged by gestational week 4-8 of human development (columns). Blue, active enhancer signature; white, no active enhancer signature. **c/d.** Left: Genomic position and evolutionary conservation of human candidate enhancer hs2656 (**c**) and its mouse ortholog mm2280 (**d**). The human sequence, but not the orthologous mouse sequence, shows evidence of H3K27ac binding at corresponding stages of craniofacial development (beige tracks). Right: Representative embryo images at e12.5 show that human enhancer hs2656, but not its mouse ortholog mm2280, drives reproducible *lacZ*-reporter expression in the developing nasal and maxillary processes at e12.5. n, reproducibility of each pattern across embryos resulting from independent transgenic integration events.

We also examined the genome-wide set of human craniofacial candidate enhancers for the presence of noncoding variants implicated in inter-individual variation in facial shape and in craniofacial birth defects through genome-wide association studies (GWAS). We aggregated lead SNPs from 41 studies of normal facial variation and craniofacial disease (**Methods**; **Supplemental Table 6**). From 1,404 lead SNPs from these studies, we identified 27,386 SNPs in linkage disequilibrium (LD; r^2^ ≥ 0.8) with the lead SNPs for the appropriate populations in the respective craniofacial GWAS. Upon intersection with H3K27ac-bound regions from bulk face tissue between stages CS13-23 (**Figure 1a**), we observed a total of 209 predicted enhancer regions overlapping 605 unique LD SNPs. This includes 43 candidate enhancer regions overlapping with 102 unique disease SNPs, and 176 candidate enhancers overlapping with 515 unique SNPs for normal facial variation (**Supplemental Table 7**).

The activity of individual enhancers can be highly dynamic across developmental stages, supporting that enhancers regulate both spatial and temporal aspects of developmental gene expression^25, 36^. To explore the temporal dynamics of human craniofacial enhancers, we determined the temporal activity profile of all 13,983 human candidate enhancers by week of development, covering gestational weeks 4 to 8 (**Figure 2b**; **Methods**). We found that a small proportion (1,624 elements or 11.6%) of elements were predicted to be continuously active as enhancers throughout all five weeks. Nearly half (6,347) showed narrow predicted activity windows limited to a single week, while another 3,137 showed continuous activity periods covering a subset of the five weeks. A smaller number of enhancers (2,236) with predicted non-continuous activities likely contains elements with truly discontinuous activity (e.g., in different subregions of the developing face), and elements not reaching significant signal at some stages, e.g., due to changes in relative abundance of cell types. In combination, these data sets provide an extensive catalog mapping the genomic location of human craniofacial enhancers, including their temporal activity patterns during critical stages of craniofacial development.

To assess the conservation of candidate enhancers identified from human tissues in the mouse model, we compared H3K27ac binding data from human developmental stages CS13-23 to previously published results for histone modifications at matched stages of mouse development^25^. The majority (12,179 of 13,983; 87%) of the human candidate enhancers are conserved to the mouse genome at the sequence level, defined by the presence of alignable orthologous sequence that is syntenic relative to surrounding protein-coding genes. Among these conserved sequences, 8,257 (59%) showed epigenomic enhancer signatures in the mouse, indicating their functional conservation. The remaining 3,922 (28%) regions were sequence-conserved but showed no evidence of enhancer activity in the mouse tissues examined (**Supplemental Table 8, Methods)**, suggesting that they are human-specific and highlighting the potential value of human tissue-derived epigenomic data for human craniofacial enhancer annotation.

To assess whether the differences in epigenomic signatures between human and mouse translate into species-specific differences in *in vivo* enhancer activity, we used a transgenic mouse assay to compare the human and mouse orthologs of a predicted human-specific enhancer. We chose a candidate enhancer located near genes *POP1*, *NIPAL2* and *KCNS2,* located in the 8q22.2 region associated with non-syndromic clefts of the face^37^ (**Figure 2c/d**). Documented mutations in *POP1* cause Anauxetic Dysplasia with pathognomonic short stature, hypoplastic midface and hypodontia along with mild intellectual disability^38–40^. We generated enhancer-*lacZ*-reporter constructs of the human and mouse orthologs of the candidate enhancer region and used CRISPR-mediated transgene insertion at the H11 safe harbor locus^41, 42^ to create transgenic mice. Embryos transgenic for the human ortholog (hs2656) show reproducible activity in the developing nasal and maxillary processes at embryonic day (e) 12.5, confirming that the human tissue-derived enhancer signature correctly predicts *in vivo* activity at the corresponding stage of mouse development (**Figure 2c**). In contrast, we did not observe reproducible craniofacial enhancer activity with the mouse orthologous sequence, concordant with the absence of enhancer chromatin marks in mouse at this location (mm2280, **Figure. 2d**).

### Transcriptome Landscape of the Mammalian Craniofacial Complex at Single-cell Resolution

To provide a higher-resolution view of the enhancer landscape of craniofacial development, we complemented these detailed maps of human craniofacial enhancers with single cell-resolved data, with the goal to identify the cell type specificity of individual enhancers. Given the genetic heterogeneity, limited availability, and processing challenges associated with early human prenatal tissues, we performed these studies on mouse tissues isolated from corresponding developmental stages (**Figure 3**).

**Figure 3.**
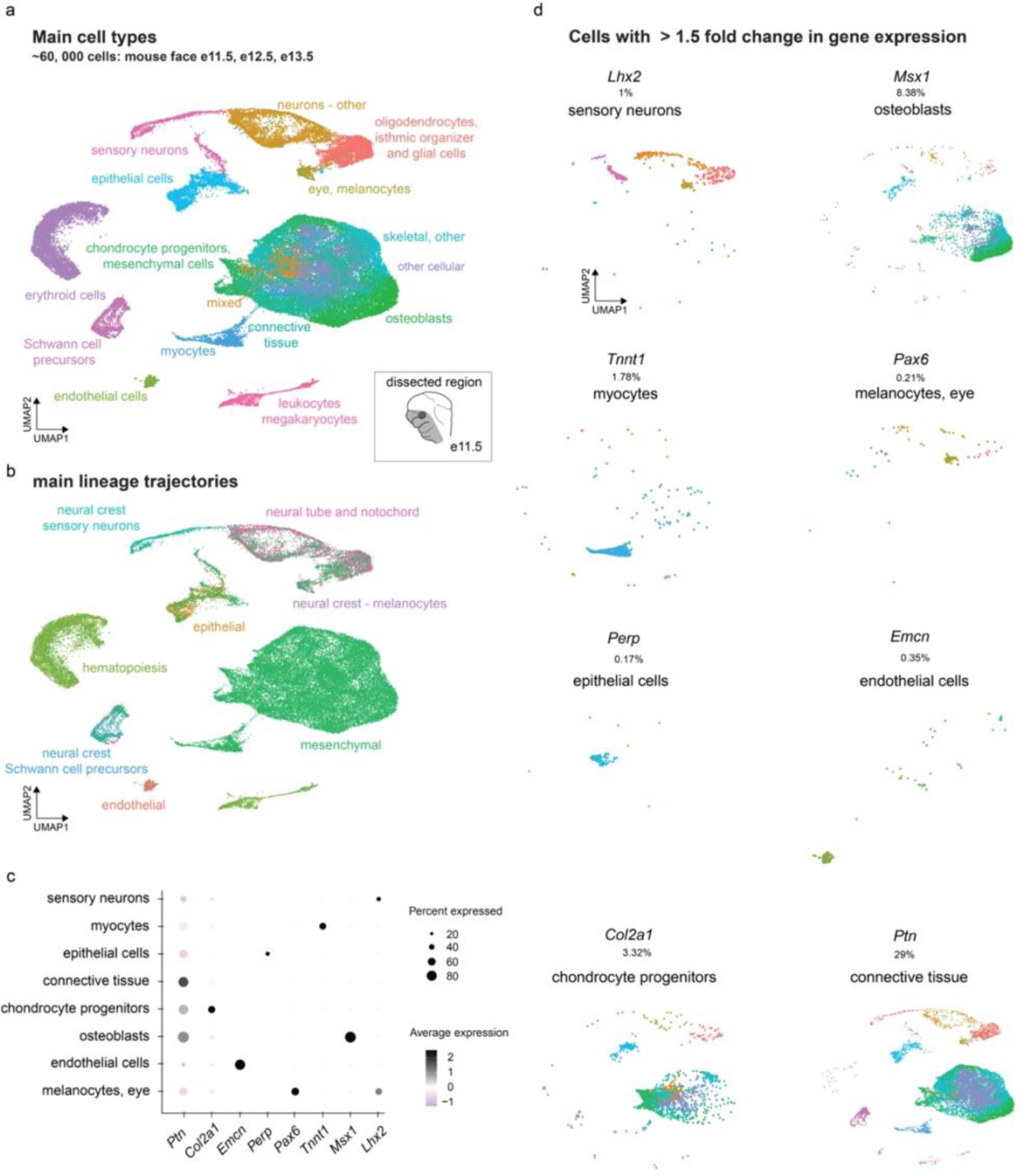
Gene expression in the mammalian craniofacial complex at single cell resolution. **a.** Uniform Manifold Approximation and Projection (UMAP) clustering, color-coded by inferred cell types across clusters from aggregated scRNA-seq for the developing mouse face at embryonic days 11.5-13.5, for 57,598 cells across all stages. Cartoon shows the outline of dissected region from the mouse embryonic face at e11.5, corresponding regions were excised at other stages. **b.** Same UMAP clustering, color-coded by main cell lineages. **c.** Expression of select marker genes in cell types shown in (a). **d.** UMAP plots comprising cells with >1.5-fold gene expression for marker genes representing specific cell types as shown in (a) and (c).

We generated a detailed transcriptome atlas from relevant stages of development and analyzed mouse facial tissue isolated from e11.5, e12.5, and e13.5 by single-cell RNA-seq (see **Methods**). Applying Uniform Manifold Approximation and Projection (UMAP) non-linear dimensionality reduction for unbiased clustering resulted in 42 primary detectable clusters (**Extended Data Figures 2-4, Supplemental Tables 9-10**). We analyzed 57,598 cells with a median of 1,659 genes expressed per cell. We systematically assigned cell type identities to the resulting clusters (**Extended Data Figures 5 and 6, Supplemental Tables 11-12**, and **Methods**) in our final <u>S</u>ingle-<u>c</u>ell <u>an</u>notated <u>Face</u> e<u>X</u>pression dataset (henceforth referred to as *ScanFaceX*), which includes 16 annotated cell types capturing the developing mammalian face and associated tissues (**Figure 3a**). Trajectory analyses using Seurat recapitulated the main lineages including epithelial, mesenchymal, endothelial, and neural crest-derived cell types including melanocytes relevant to face development (**Figure 3b**). The final annotated cell type clusters showed strong cluster-specific expression of established markers genes relevant to craniofacial development such as *Col2a1* (chondrocyte progenitors)^43–45^, *Msx1* (osteoblasts)^46–48^, *Perp* (epithelial cells)^49, 50^, *Emcn* (endothelial cells)^51, 52^, *Lhx2* (sensory neurons)^53, 54^, *Pax6* (melanocytes)^55, 56^, *Tnnt1* (myocytes)^57^, and *Ptn* (connective tissue)^58^ (**Figure 3c and 3d, Extended Data Figure 7**). These benchmarking results indicate that *ScanFaceX* provides an accurate single-cell transcriptome reference for relevant stages of craniofacial development that can serve as a foundation for integration with other chromatin data types.

**Figure 4.**
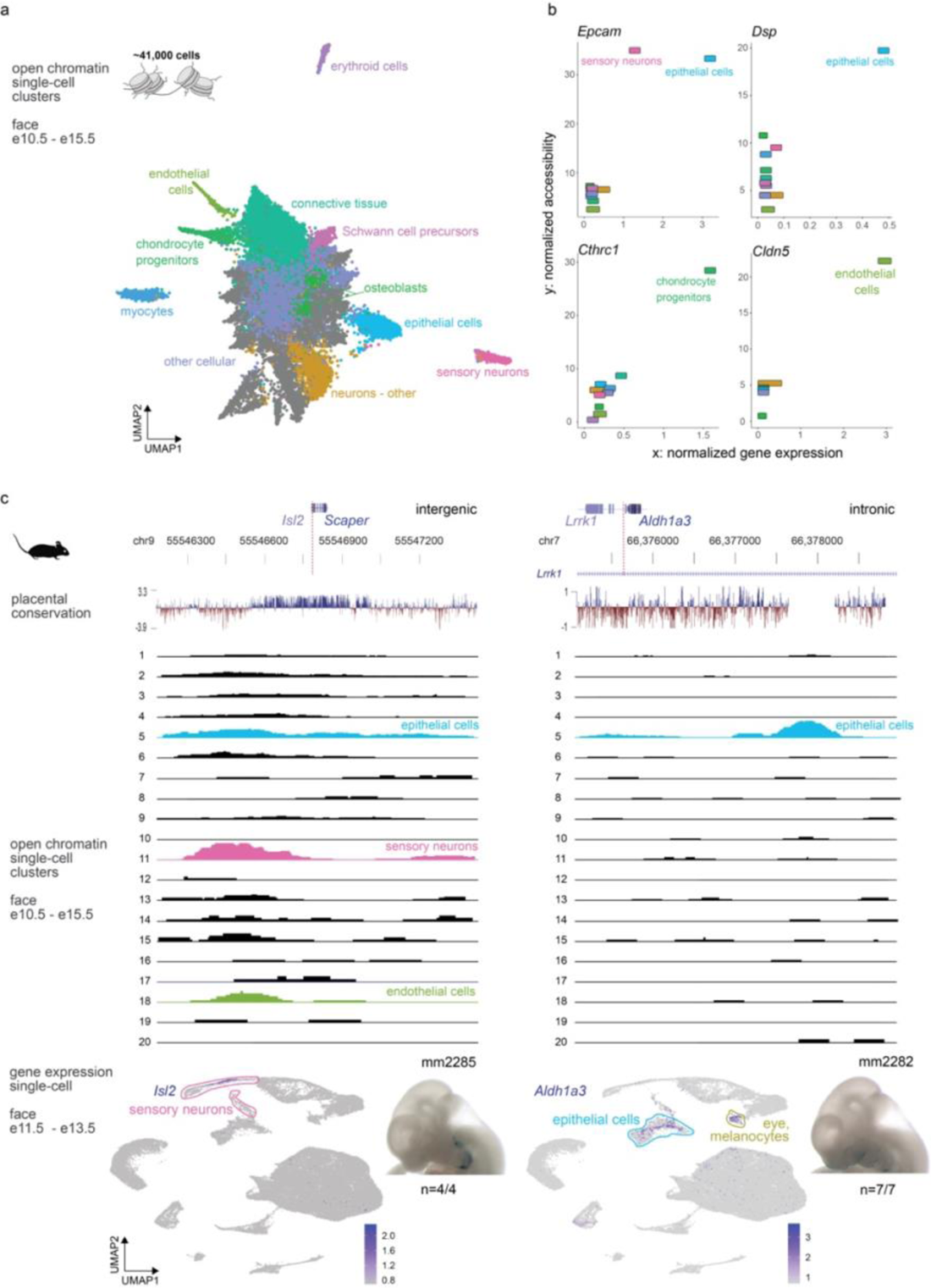
Differential chromatin accessibility at craniofacial *in vivo* enhancers correlates with cell type-specific expression of nearby genes. **a.** Unbiased clustering (UMAP) of open chromatin regions from snATAC-seq of the developing mouse face for stages e10.5-15.5 for approximately 41,000 cells. The cell types are assigned based on label transfer (Seurat) from cell-type annotations of the *ScanFaceX* data. **b**. Correlation between normalized gene expression (x-axis) from *ScanFaceX* and normalized accessibility (y-axis) from snATAC-seq for select genes (*Epcam, Dsp, Cthrc1, Cldn5*) and their transcription start sites with the highest correlation evident in relevant cell types. **c**. Genomic context and evolutionary conservation (in placentals) for corresponding regulatory regions in the vicinity of the *Isl2/Scaper* locus, and an intronic distal enhancer within *Lrrk1.* Tracks for individual snATAC-seq clusters from developing mouse face tissue (e10.5 to e15.5), with cluster-specific open chromatin signatures for relevant annotated cell types are shown for the same genomic regions. UMAP of *ScanFaceX* data shows expression of *Isl2* and *Aldh1a3* (gene adjacent to *Lrrk1*) in expected cell-types. Images for a representative mouse embryo at e11.5 for both loci show validated *in vivo lac-Z*-reporter activity of the respective regions. n, reproducibility of each pattern across embryos resulting from independent transgenic integration events.

**Figure 5.**
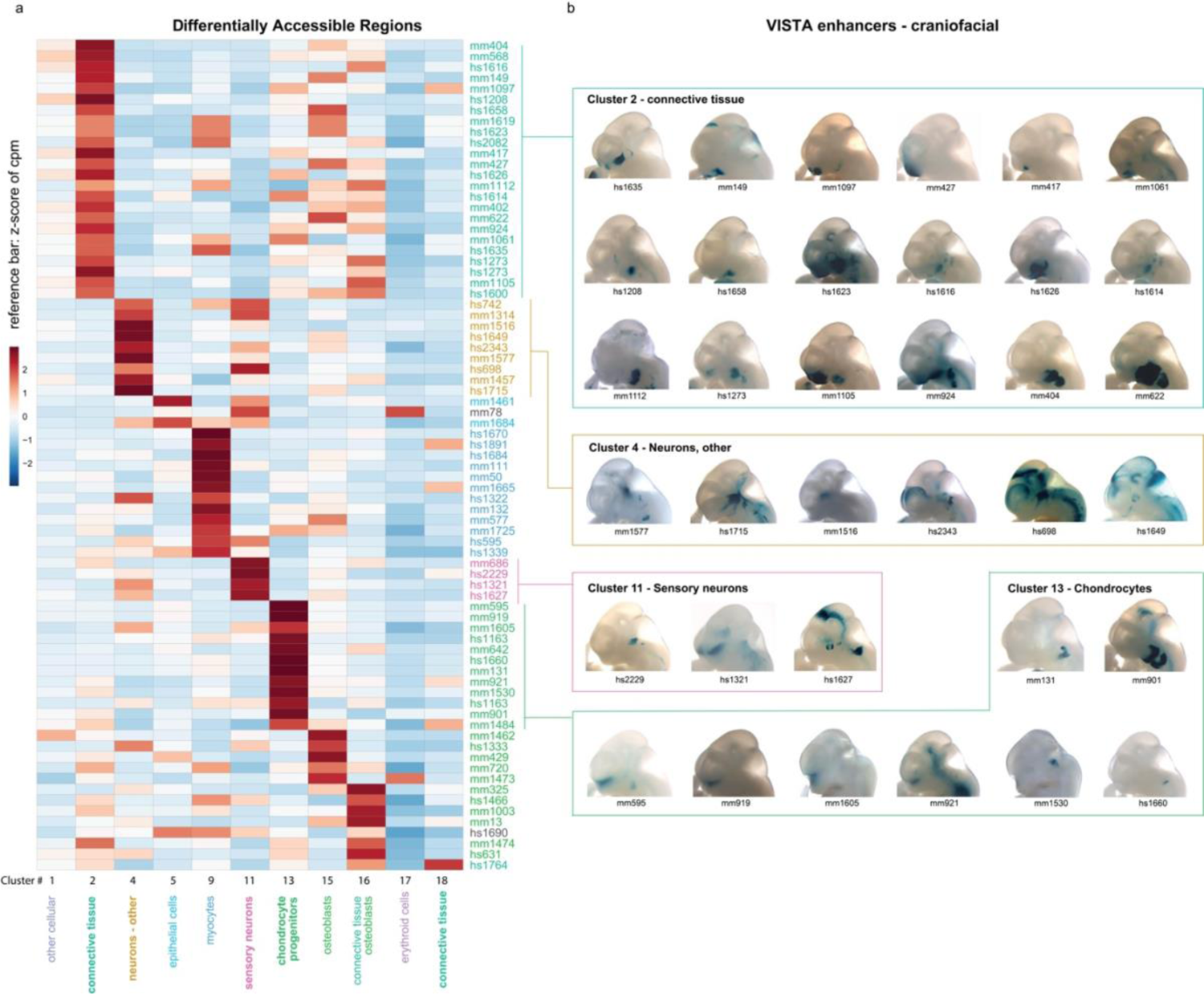
Cell type-specific chromatin accessibility of craniofacial *in vivo* enhancers. **a.** Heatmap indicates the chromatin accessibility of 77 craniofacial *in vivo* enhancers in 11 major cell type clusters. cpm: counts per million. **b.** Representative images of transgenic embryos from VISTA Enhancer Browser, showing *in vivo* activity pattern of 35 selected enhancers at e11.5. Embryo images are grouped by example cluster-types from (a) in this retrospective assignment.

**Figure 6.**
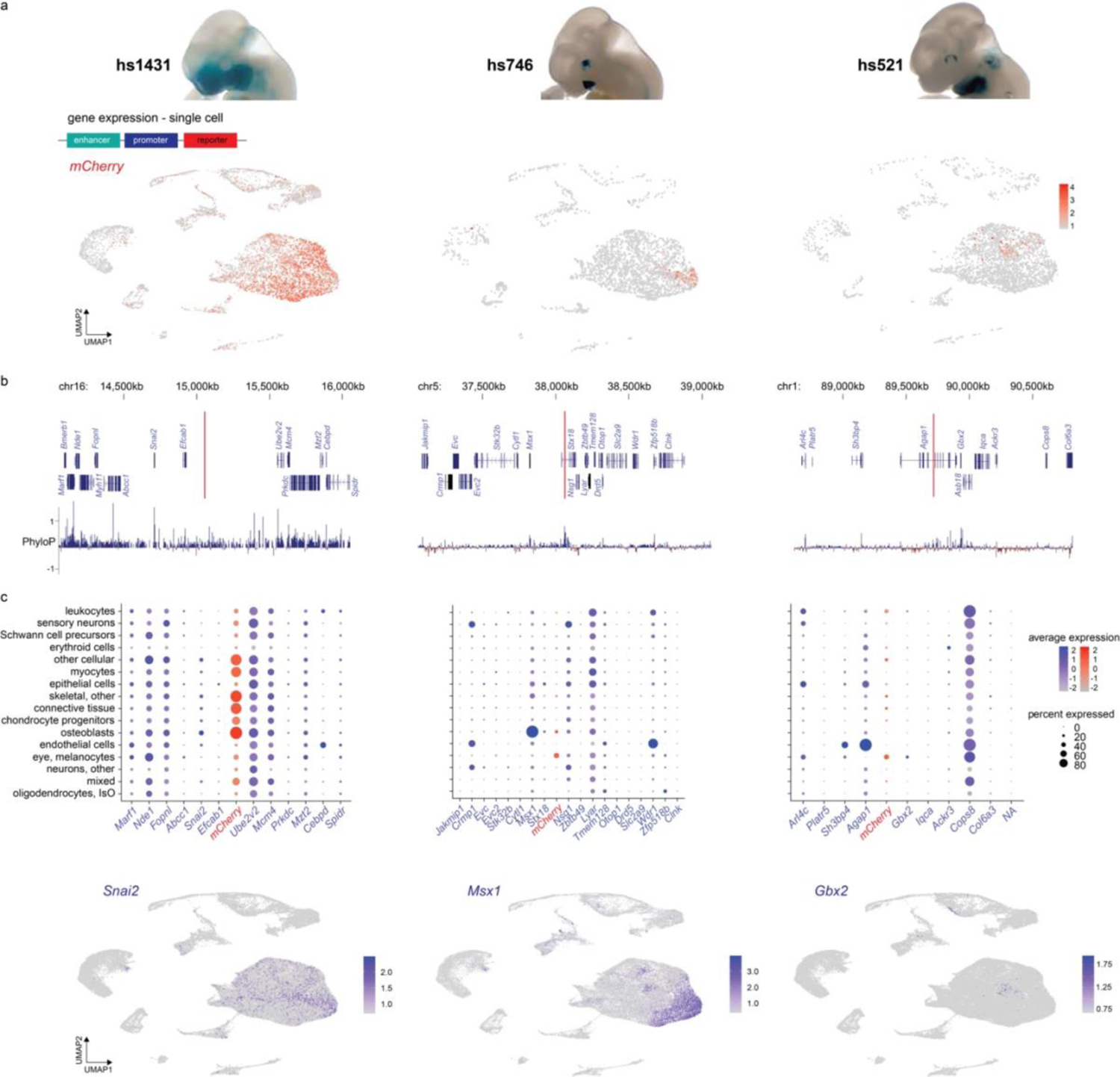
Cell type-specific enhancer activity at single-cell resolution. **a.** *in vivo* activity pattern of select craniofacial enhancers (hs1431, hs746, hs521) at e11.5, visualized by *lacZ*-reporter assays (top). In separate experiments, the same enhancers were coupled to an *mCherry*-fluorescent reporter gene and examined by scRNA-seq of craniofacial tissues of resulting embryos. UMAPs show enhancer-driven *mCherry* expression (see **Fig. 3a** for reference). **b.** Location of enhancers hs1431, hs746 and hs521 in their respective genomic context (red vertical lines), along with protein-coding genes within the genomic regions and local conservation profile (PhyloP). **c.** Average expression of genes (Seurat) in the vicinity of the respective enhancers, and proportion (percent) of cells expressing the genes in specific cell types. Enhancer-driven *mCherry* signal is plotted in the center *in lieu* of the approximate enhancer location in its endogenous genomic context. Bottom panels show expression of *Snai2*, *Msx1,* and *Gbx2* as likely candidate target genes for each of the enhancers hs1431, hs746 and hs521 across UMAPs. IsO: Isthmic Organizer Cells.

### Differential Chromatin Accessibility Correlates with Cell Type-specific Signatures

To identify developmental enhancers at single-cell resolution, we performed single-nucleus ATAC-seq (snATAC-seq)^59^ on mouse face embryonic tissues at select developmental time points (**Figure 4**). Across all stages analyzed, 41,483 cells that passed all quality control steps were considered in the final analysis, and their unbiased clustering resulted in 20 discernable clusters (see **Methods**). Out of a total of 115,521 open chromatin regions in the snATAC-seq data, we observed 16,564 differential accessible regions (DARs) across 20 separate clusters, indicating that each of the clusters representing identical or very similar cell types have distinct open chromatin signatures (**Extended Data Figure 8, Supplemental Table 13**). Next, we integrated our single-cell open chromatin data with the cell type annotations from *ScanFaceX* single-cell transcriptome data using Seurat-based label transfer (see **Methods**). Upon integration, a substantial subset of DARs (10,038 out of 16,564; 60%) were retained, and developing craniofacial cell types including chondrocytes, osteoblasts, myocytes and connective tissue, epithelial cells, melanocytes, and sensory neurons showed high correlation between the two data types (**Figure 4a-b, Extended Data Figures 9 and 10; Methods**). Chromatin accessibility at putative distal enhancer regions as well as transcription start sites showed distinct cell type specificity. For example, the representative intergenic region near *Isl2* and *Scaper,* and an intronic region of *Lrrk1* differentially active in clusters representing sensory neurons and/or epithelial cells, illustrate the resolution of our data relative to previously available predictions from bulk face tissue^25, 60, 61^ (**Figure 4c**). These predictions of cell type specificity are well aligned with known functions of the respective target genes. *Isl2* has been shown to be selectively expressed in a subset of retinal ganglion cell axons that have important functions in binocular vision^62^. Allelic variants and mutations in *SCAPER* cause intellectual disability with retinitis pigmentosa in humans^63–65^. The *Lrrk1* intronic element is near *Aldh1a3,* a gene adjacent to *Lrrk1;* mutations in the orthologous *human ALDH1A3* cause an autosomal recessive form of isolated microphthalmia^66–69^. These putative enhancer regions near *Isl2* and *Scaper*, and in the intron of *Lrrk1* drive reproducible *lacZ*-reporter activity in the developing mouse face in anatomical regions that are consistent with neuronal and epithelial cell types (**Figure 4c)**. Both *Isl2* and *Aldh1a3* are highly expressed in sensory neurons and epithelial cell clusters, respectively, in *ScanFaceX* data (**Figure 4c**). In an additional example, an enhancer near the promoter region of *Mymx*, which is exclusively active in the myocyte cluster, coincides with *Mymx* expression in myocytes in *ScanFaceX* (**Extended Data Figure 11**).

To facilitate utilization of the full set of genome-wide, cell type-resolved enhancer predictions, we used these mouse tissue-derived single-cell enhancer predictions in combination with our human bulk tissue-derived enhancer catalog, to generate a <u>S</u>ingle-<u>c</u>ell <u>an</u>notated <u>Face</u> e<u>N</u>hancer (*ScanFaceN*) catalog of human enhancer regions with predicted activity profiles across craniofacial cell types (**Supplemental Tables 14-16**). The majority (7,899 of 13,983; 56%) of human tissue-derived facial candidate enhancers overlap with an accessible chromatin region in at least one cluster of our *ScanFaceN* catalog, and 2,339 (30%) of these regions overlap with DARs in *ScanFaceN*.

### Cell Type-resolved Enhancer Predictions and *in vivo* Enhancer Activity Patterns

To explore the relationship between predicted cell type specificities of enhancers and their respective spatial *in vivo* activity pattern during craniofacial development, we intersected the *ScanFaceN* DARs from the 11 main *ScanFaceX*-matched clusters with craniofacial enhancers validated *in vivo* and curated in the VISTA Enhancer Browser^34^ (**Figure 5a**). We observed general correlations between cluster-specific accessibility and spatial *in vivo* patterns among 77 enhancers that showed chromatin accessibility in at least one of the 11 main clusters. For example, the connective tissue-mesenchymal cluster (cluster 2) of the craniofacial snATAC-seq tends to group VISTA enhancers with activity specific to the branchial arches, while the chondrocyte cluster (cluster 13) has multiple VISTA enhancers with activity in the mid-face, paranasal regions, and/or the region at the junction of the developing forebrain and nasal prominences that are consistent with developing cartilaginous regions of the face (**Figure 5b**). Despite these broad correlations, we observed considerable heterogeneity of spatial patterns within most clusters, underscoring the spatiotemporal complexity of craniofacial morphogenesis, which relies on cell type-specific regulatory programs in combination with highly regionalized regulatory cues.

### Cell Type-specific Enhancer Activity at Single-cell Resolution

To explore whether craniofacial enhancer activity can be quantitatively assigned to specific cell types *in vivo*, we generated transgenic mice in which selected craniofacial enhancers were coupled to a fluorescent *mCherry* reporter gene (**Figure 6a**). We examined three different craniofacial enhancers (hs1431, hs746 and hs521), two of which (hs1431 and hs746) we previously demonstrated to be required for normal facial development^1^ (**Figure 6b**). In all cases, we isolated craniofacial tissue from transgenic reporter embryos at e11.5 and performed scRNA-seq (**Figure 6a**). For hs1431, near *Snai2*, which is active across many regions of the developing face, *mCherry* expression is observed across almost all cell type clusters, indicating that hs1431 is broadly active across multiple cell types during craniofacial development (**Figure 6c**). In contrast, hs746 which is in the vicinity of *Msx1*, is primarily active in osteoblasts and in a subset of cells expressing *Msx1*, a gene previously shown to regulate the osteogenic lineage^70^. Enhancer hs521, located near *Gbx2*, is primarily active in a subset of mesenchymal cells and chondrocyte progenitors, and its activity coincides with a subset of cells expressing *Gbx2* (**Figure 6c**), a gene known to be active in the developing mandibular arches^9^. Together, these data illustrate how purpose-engineered enhancer-reporter mice can be used to validate and further explore the *in vivo* activity patterns of craniofacial enhancers identified through genome-wide single-cell profiling studies.

## Discussion

The lack of data from primary tissues and incomplete mapping of human developmental enhancers in craniofacial morphogenesis has been a challenge in the systematic assessment of the role of enhancers in craniofacial development and disease. In the present study, we have generated human bulk and mouse single-cell data to create a comprehensive compendium of enhancers in human and mouse development, including temporal profiles and predictions of cell type specificity. We identify major cell populations of the developing mammalian face, along with corresponding genome-wide enhancer profiles. We also show that while many enhancers are functionally conserved between human and mouse, additional human-specific enhancers that show no functional conservation in mice can be identified by profiling human tissues. Our data illustrate the considerable temporal dynamics of human craniofacial enhancers, a critical aspect for understanding the developmental timing of enhancer activity related to specific phenotypes such as clefts and mid-facial deformities. As clinical sequencing becomes increasingly common and accessible to both patients and the medical community, our data may serve as an essential resource to address the gaps in understanding the potential pathogenicity of regulatory variants.

The single-cell resources generated through this study, *ScanFaceX* for expression and *ScanFaceN* for enhancers, contain a total of 115,521 candidate enhancers as defined by chromatin accessibility, including 10,038 that show differential chromatin accessibility across annotated cell types in face morphogenesis. We demonstrated how engineered mice can be used to study these enhancers *in vivo* at single-cell resolution. Using a transgenic reporter assay coupled to single-cell RNA-seq, we defined the cell type-specific activity of three craniofacial enhancers during embryonic development. This approach illustrates how these methods can be combined to determine the *in vivo* cell type specificity of individual enhancers and relate their activity to cell type-specific expression of their putative target genes. All of these data are also available in FaceBase and the VISTA Enhancer Browser for community use^1,61, 71^. In summary, our work provides a multifaceted and expansive resource for studies of craniofacial enhancers in human development and disease.

## Acknowledgements

This work was supported by U.S. National Institutes of Health (NIH) grant to A.V (R01DE028599 and U01DE024427). Research was conducted at the E.O. Lawrence Berkeley National Laboratory and performed under U.S. Department of Energy Contract DE-AC02-05CH11231, University of California (UC). The human embryonic face samples were provided by the Joint MRC / Wellcome Trust (MR/R006237/1, MR/X008304/1 and 226202/Z/22/Z) Human Developmental Biology Resource^72^ (www.hdbr.org). F.D. was supported by the Swiss National Science Foundation grant P400PB_194334.

## Author Contributions

S.S.R., D.E.D., L.A.P., and A.V. designed the study. S.S.R., C.S., M.O., Y.Zhu, H.W., S.Y.A., J.A.A., V.A., S.T., I.P-F., C.S.N., M.Kato., R.H., K.V.M., A.W., L. L., S.P. performed experiments. J. A. A. performed imaging. S.S.R., K.P., M.L.A., M. Kosicki, L.E.C., F.D., M.B., G.K., I.B., Y.F-Y. analyzed data. S.S.R., L.P., and A.V. wrote the manuscript with input from the remaining authors. We thank Yoon Gi “Justin” Choi of the University of California, Berkeley QB3 Genomics Core for technical assistance with the 10X Genomics set up.

## Methods

### Human Subjects

All aspects of this study involving human tissue samples were reviewed and approved by the Human Subjects Committee at Lawrence Berkeley National Laboratory (Protocol Nos. 00023126 and 00022756). Human embryonic face samples were obtained from the Human Developmental Biology Resource at Newcastle University (HDBR, hdbr.org), in compliance with applicable state and federal laws and with fully informed consent. No identifying information for human samples was shared by HDBR. All embryonic samples for which primary data was generated in this study were verified to be karyotypically normal by HDBR. Primary data from embryonic whole face samples at post-conception weeks 7 and 8 were generated for this study. All embryonic samples were shipped on dry ice and stored at −80°C until processed. ChIP-seq data for three samples at Carnegie stage (CS)18, one sample at CS 19, two samples at CS22 and one sample at CS23 are presented in this study, along with accompanying ATAC-seq data for two samples at CS18, one sample at CS19, one sample each at CS22 and CS23. Processed data for CS 13-16 was obtained from previously published studies ^26^ and included in our downstream integrative analyses. All datasets are listed in **Supplemental Table 1**.

### Animal Studies and Experimental Design *in vivo*

All animal work was reviewed and approved by the Lawrence Berkeley National Laboratory Animal Welfare Committee. Mice used for this study were housed at the LBNL Animal Care Facility, which is fully accredited by AAALAC International. All mice were routinely health checked and monitored daily for food and water intake. Mice (*Mus musculus*; FVB strain) across developmental stages from embryonic day 10.5 through 15.5 were used in this study. Animals of both sexes were used in the analysis. Sample size selection and randomization strategies were followed based on our previous experience of performing transgenic mouse assays for ∼3000 published enhancer candidates^41, 42^. Mouse embryos that lacked the reporter transgene or were not at the correct developmental stage were not included further in analysis.

### Transgenic Mouse *Assays in vivo*

Transgenic enhancer-reporter assays were performed as previously described ^41, 42^. Briefly, a minimal *Shh* promoter and reporter gene were integrated into a non-endogenous, safe harbor locus ^42^ in a site-directed transgenic mouse assay. The selected genomic region was PCR amplified from human or mouse genomic DNA where applicable; the PCR amplicon was cloned into a *lacZ*-reporter vector (Addgene #139098) or *mCherry* reporter vector (available upon request) using Gibson assembly (New England Biolabs) ^73^. The final transgenic vector consists of the predicted enhancer–promoter–reporter sequence flanked by homology arms intended for the *H11* locus in the mouse genome. Sequence of the cloned constructs was confirmed with Sanger sequencing or MiSeq. Transgenic mice were generated using pronuclear injection, as previously described^42^. Embryos were collected for various downstream experiments at embryonic days 10.5 through 15.5. Beta-galactosidase staining was performed in our standardized pipeline with the following modification. Embryos were fixed with 4% paraformaldehyde (PFA) for 30 minutes for E11.5 embryos, respectively, while rolling at room temperature. The embryos were genotyped for presence of the transgenic construct. Embryos positive for transgene integration into the *H11* locus and at the correct developmental stage were considered for comparative reporter gene activity across respective stages and were imaged on a Leica MZ16 microscope. Genomic coordinates for VISTA enhancer hs2656 (**Figure 2**); enhancer mm2280 (**Figure 2**), mm2282 and mm2285 (**Figure 4**), and mm2281 (**Extended Data Figure 11**) are shown in **Supplemental Table 5** and **17** respectively.

For transgenic experiments demonstrating cell type-specific enhancer activity at single-cell resolution and involving hs1431, hs746 and hs521 (**Figure 6**), a combination of *Hsp68* promoter and *mCherry* reporter were used.

### ChIP-seq

Chromatin immuno-precipitations were performed as previously described ^74^. Briefly, frozen and non-cross-linked face tissue was dissociated in PBS by pipetting until homogenized and cross-linked with 1% formaldehyde at room temperature. Cells were lyzed and chromatin was sonicated using a Biorupto device (Diagenode) to obtain fragments with an average size ranging between 100–600 bp. Input sample was set aside and stored appropriately, Protein A and G Dynabeads (Invitrogen) were added to the sample, and chromatin was incubated for 2h at 4°C with 5 μg of anti-H3K27ac antibody (Active Motif, Cat# 39133, Lot 01613007). Immuno-complexes were sequentially washed, and the immunoprecipitated DNA complexes were eluted in an SDS buffer at 37°C for one hour. Samples were reverse-crosslinked with with Proteinase K overnight at 37°C. DNA was purified with a ChIP DNA clean concentrator (D5205 Zymo Research), and a KAPA SYBR Green qPCR mix was used to assess presence of H3K27 acetylated regions versus negative control regions. DNA was quantified using Qubit, and size distribution and DNA concentration of the samples were assessed on the Agilent Bioanalyzer. Illumina TruSeq library preparation kit was used for downstream library preparation, and libraries were sequenced as single-end 50 bp reads on an Illumina HiSeq 2500.

ChIP-seq data was analyzed using the ENCODE histone ChIP-seq Unary Control Unreplicated pipeline (https://www.encodeproject.org/pipelines/ENCPL841HGV/) implemented at DNAnexus (https://www.dnanexus.com). Briefly, reads were mapped to the human reference genome version hg38 using BWA (v0.7.7) and sorted bam file generated using samtools (v0.1.19). Peak calling was performed using MACS2 (v2.2.4; --broad flag, q-value < 0.05), and overlapping peaks identified using overlap_peaks.py. A combined peak set was called by merging peaks from all samples. Merged peaks within 1kb of transcription starts sites as defined by GENCODE were removed, resulting in 70,075 distal peaks. Of those, 13,983 peaks were present in at least two samples in each embryonic week which were retained for final analysis.

### ATAC-seq

Embryonic samples were processed for ATAC-seq as previously described ^74^. In short, harvested tissues were lysed, centrifuged for 10min at 500 x g, at 4°C, and the resulting cell pellet was treated with the Nextera DNA transposase Tagment DNA Enzyme (Catalog number: 20018705) and the transposed DNA was eluted using Qiagen MinElute PCR purification kit. Samples were then PCR amplified using the NEB Next High-Fidelity 2xPCR Master Mix (catalog number: NEBE6040SEA) with Nextera PCR primers 1 (AATGATACGGCGACCACCGAGATCTACACNNNNNNNNTCGTCGGCAGCGTC) and 2 (CAAGCAGAAGACGGCATACGAGATNNNNNNNNGTCTCGTGGGCTCGG), and DNA was purified as described above. The eluted library was analyzed for quality in a Bioanalyzer High Sensitivity assay and samples were subsequently deep sequenced on an Illumina HiSeq2500. ATAC-seq data was analyzed using the ENCODE ATAC-seq (unreplicated) pipeline (https://www.encodeproject.org/pipelines/ENCPL344QWT/).

### RNA-seq

Samples were processed for RNA-seq and libraries were generated as previously described^74, 75^. Briefly, RNA was isolated from the dissociated face tissue using TRIzol Reagent (Life Technologies), all samples were DNase-treated (TURBO DNA-free Kit, Life Technologies), and assessed for quality (RNA 6000 Nano Kit, Agilent) on a 2100 Agilent Bioanalyzer. TruSeq Stranded Total RNA with Ribo-Zero Human/Mouse/Rat kit (Illumina) was used to prepare RNA-seq libraries according to manufacturer’s protocol. RNA-seq libraries were depleted of high molecular weight products in an Illumina Resuspension Buffer and by incubating in 60 μL Agencourt AMPure XP beads for 4 min. AMPure beads were pelleted, washed twice with 80% ethanol and the DNA was eluted per manufacturer’s instructions. RNA concentration and quality of the RNAseq libraries were assessed using a 2100 Bioanalyzer with the High Sensitivity DNA Kit (Agilent), and libraries were sequenced as single-end 50 bp reads on an Illumina HiSeq 2500.

RNA-seq data was analyzed using the ENCODE RNA-Seq (Long) Pipeline-1 replicate pipeline (https://www.encodeproject.org/pipelines/ENCPL002LSE/) implemented at DNAnexus (https://www.dnanexus.com). Briefly, reads were mapped to the reference genome using STAR align (V2.12). Genome wide coverage plots were generated using bam to signals (v2.2.1). Gene expression counts were generated using RSEM (v1.4.1). Human datasets were analyzed using human reference genome version hg19, and GENCODE v24 gene annotations. Mouse datasets were analyzed using mouse reference genome version mm10 and GENCODE M4 gene annotations.

### GWAS Data

The NHGRI-EBI Catalog of Genome-wide association studies^76^ was mined for studies with the following keywords: craniofacial, face, cleft lip, cleft palate, microsomia, salivary, taste, and tooth. The compiled studies comprised of diverse populations and ethnicities ranging from those belonging to the Unites States, Europe, Taiwan, China, Singapore, Korea and the Philippines, Brazil, Spain, Latin Americas, Uyghurs as well as admixed populations. For data published in the catalog by early 2022, we aggregated 41 studies representing normal facial variation as well as dento-oro-craniofacial disease. The SNiPA tool^77^ was used for querying SNPs in linkage disequilibrium (r^2^ ≥ 0.8) with the lead SNPs for the appropriate populations for the respective GWAS. This compilation of GWAS (**Supplemental Tables 6-7**) was intersected with 13,983 highly reproducible human enhancers derived from primary embryonic bulk face between CS13-23.

### Intersecting VISTA Catalog with Predicted Craniofacial Enhancers

We intersected the 3,193 curated enhancers in the VISTA Enhancer Browser with 13,983 reproducible human candidate enhancers from this study or 10,038 mouse DARs, requiring a minimum 100bp overlap. We used liftOver to obtain the genomic coordinates in species relevant assembly (**Extended Data Figure 1**, and **Supplemental Table 4**).

### Single-cell RNA-seq

Transgenic embryos were harvested at the determined developmental stage, between 11.5 - 13.5 dpc (8 samples at e11.5, 1 sample at e12.5, and 4 samples at e13.5), and examined for positive *mCherry* signal. Embryos positive for *mCherry* reporter activity showed reproducible and comparable enhancer-reporter expression as seen in the *lacZ* expression patterns for VISTA enhancers hs1431, hs521 and hs746 used in this study. Embryos were consistently kept in ice-cold PBS until dissection. Upon fluorescent screening, developing face tissue was dissected with the aid of a Leica MZ16 microscope, and immediately processed for downstream experiments. Fresh mouse embryonic face tissue was mechanically dissociated by pipetting gently into a single-cell suspension using Accumax, assessed for viability of cells and cell density using Trypan Blue staining. Individual cells were quantified, spiked with 10% HEK293T/17 frozen-thawed cells, and processed using the 10X Genomics Chromium Next GEM Single Cell 3’ protocol including transcript capture and library preparation for single-cell gene expression. Samples were either processed individually or pooled using a previously described Multi-seq strategy^78^ upstream of the 10X Genomics Chromium protocol. The resulting libraries were sequenced on an Illumina HiSeq2500 or NovaSeq 10X. BCL files from Illumina were processed into FASTQ format, individual sample libraries were de-multiplexed as necessary, reads were aligned to mm10 reference genome where *mCherry* sequence was added as an additional chromosome. Cell Ranger 3.1.0 software was used to process the raw sequence files and generate feature-barcode matrices. After correcting for batch effects, data from all libraries was aggregated into a single R object file using the 10X Genomics Cell Ranger 3.1.0. Seurat v3.2 guided clustering tutorial was used for formal downstream analyses ^79–81^. Adhering to the standard pre-processing workflow and quality control, cells with unique feature counts between 200-2000 and < 5%mitochondrial reads were retained. Normalization, feature selection, scaling, dimensional reduction, clustering and finding cluster biomarkers i.e., differentially expressed features were performed as guided. Our final Seurat/clustered UMAP consists of a 25,645 feature by 57,598 cell matrix.

#### Assigning cell-type identity to scRNA-seq clusters

We systematically assigned cell type identities to the clusters in our craniofacial scRNA-seq dataset using two computational methods. (i) Using our primary single cell dataset as query, we assigned cell type identities by Seurat-based automated reference mapping to a previously published large single-cell gene expression dataset ^82^ of whole mouse embryonic development for stages e9.5-13.5, the reference was down sampled to 100K cells for efficient processing and retained all 38 broad cell types originally described. 27 cell types from the reference were summarily mapped in our craniofacial scRNA-seq dataset by Seurat’s label transfer; the referenced cell types showed a good overall correlation with the cell types associated with the top 20 marker genes in most clusters in our *ScanFaceX* dataset. (ii) In parallel, we used the scoreMarkers wrapper function previously described in the *scran* package which uses effect sizes (Cohen’s *d* statistic) to perform differential expression to list marker genes for each of the clusters in a scRNA-seq dataset ^83^. These marker gene sets were tested for enrichment of Gene Ontology (GO) biological process terms by performing a hypergeometric test to identify GO terms overrepresented in our *ScanFaceX* dataset. Cell-type annotations from methods (i) and (ii) described above were compared and resulted in each cluster in the *ScanFaceX* dataset having one or more cell-type annotations. Finally, cell clusters that showed similar or close cell-type specific signatures were manually merged to reflect 16 formal annotations for definitive cell types capturing craniofacial development and morphology (**Supplemental Tables 11-12**).

#### Single-nucleus ATAC-seq

Face tissue was dissected from mouse embryos for each of the developmental stages e10.5-15.5, and flash frozen and stored at −80°C until ready to process. Tissue was transported to the Center for Epigenomics, University of California, San Diego School of Medicine, La Jolla, CA for processing using a combinatorial indexing-assisted single nucleus ATAC-seq previously described ^59^. Briefly, nuclei were isolated and permeabilized in optimized conditions, pelleted and suspended in resuspended in 500μL high salt tagmentation buffer. Nuclei were counted using a hemocytometer and 2,000 nuclei were dispensed into each well of a 96-well plate per sample. A BenchSmart™ 96 (Mettler Toledo) was used to add 1μL barcoded Tn5 transposomes to each of the wells in the 96-well plate, the mix was incubated for 60 min at 37 °C with shaking (500 rpm). EDTA at a final concentration of 20mM was then added to each well for incubation at 37 °C for 15 min with shaking (500 rpm) to terminate the Tn5 reaction. Next, nuclei were suspended in 20 μL of 2x sorting buffer (2 % BSA, 2 mM EDTA in PBS), wells for each sample were combined and stained with Draq7 at 1:150 dilution (Cell Signaling). 20 nuclei per sample were sorted per well into eight 96-well plates (total of 768 wells) in 10.5 μL of Elution Buffer (25 pmol primer i7, 25 pmol primer i5, 200 ng BSA (Sigma) using a Sony SH800. A Biomek i7 Automated Workstation (Beckman Coulter) was used for performing downstream steps. Samples were incubated at 55 °C for 7 min with shaking (500 rpm) in 1 μL 0.2% SDS, followed by addition of 12.5% Triton-X to quench the SDS. Samples were PCR-amplified (12.5 μL NEBNext High-Fidelity 2× NEB PCR Master Mix; [72 °C 5 min, 98 °C 30 s, (98 °C 10 s, 63 °C 30 s, 72°C 60 s) × 12 cycles, held at 12 °C]). Wells were combined post-PCR. A manual MinElute PCR Purification Kit (Qiagen) along with a vacuum manifold (QIAvac 24 plus, Qiagen) was used for library purification, and size selection was performed with SPRISelect reagent (Beckmann Coulter, 0.55x and 1.5x). A Qubit fluorimeter (Life Technologies) was used to quantify the libraries and the nucleosomal pattern of fragment size distribution was verified on a High Sensitivity D1000 Tapestation (Agilent). Libraries were sequenced on a NextSeq500 or HiSeq4000 (Illumina) using custom sequencing primers.

Reads were aligned to mm10 reference genome using bowtie2 with default parameters and cell barcodes were added as a BX tag in the bam file. Only primary alignments were kept. Duplicated read pairs were removed with Picard, and proper read pairs with insert size less than 2000 were kept for further analysis.

#### Clustering and cell-type annotation

snapATAC package was used to perform read counting and cell clustering for both all-tissue clustering and tissue-level clustering ^84^. First, we removed nuclei with less than 400 fragments or TSS enrichment < 4 for all tissues and calculated a cell-by-bin matrix at 5000-bp resolution for every sample independently, binarized the matrices and subsequently merged them for each clustering task. Next, we filtered out any bins overlapping with ENCODE blacklist (mm10, http://mitra.stanford.edu/kundaje/akundaje/release/blacklists/mm10-mouse/mm10.blacklist.bed.gz). We then normalized the read coverage of all bins with log10 (count+1) and Z-score transformation, and only removed bins with absolute Z scores higher than 2. After these filtering steps, we calculated Jaccard Index and performed dimensional reduction using the runDiffusionMaps function on similarity matrices. The memory usage of the matrices scales quadratically with the number of nuclei. Therefore, we sampled a subset of 30,000 “landmark” nuclei to compute the matrices and then extended to the rest of the cells. After dimensional reduction, we selected top 20 eigenvectors based on the variance explained by each eigenvector and computed 20 nearest neighbors for each nucleus and applied the Leiden algorithm to define 20 clusters.

To perform label transfer from the scRNA-seq to the corresponding snATAC-seq data we first created a gene activity matrix from the snATAC-seq data using accessibility in TSS and gene bodies with the SnapATAC package. We then converted our gene activity matrix into a Seurat object and used default parameters for the Seurat function FindTransferAnchors to perform canonical correlation analysis on the gene activity matrix along with the gene expression quantification from the scRNA-seq data. Finally, we used the TransferData function to annotate the snATAC-seq data via label transfer.

For the scatter plots showing normalized accessibility versus gene expression (**Figure 4b**), we used a gene by cell matrix which has counts for reads at the TSS and the gene body of each gene. To explore the accessibility for VISTA elements hs1431 and hs746, we generated UMAP plots using accessibility at the corresponding open chromatin region (overlapping peak call). For hs521, we assigned accessibility at the nearest peak as there was no overlapping open chromatin region called. The plotted colors are scaled by RPM and smoothed.

## Statistical Analyses

Statistical analyses are described in detail in the Methods sections above. Whenever a p-value is reported in the text, the statistical test is also indicated. All statistics were estimated, and plots were generated using the statistical computing environment R (www.r-project.org).

## Imaging

For both brightfield and fluorescent images, all embryos were imaged with a Leica MZ16 microscope and a Leica DFC420 digital camera using identical lighting conditions.

## Data Availability

The ChIP-seq, ATAC-seq, RNA-seq as well as scRNA-seq and snATAC-seq data presented in this publication, and generated as part of this study are accessible at the National Institute of Dental and Craniofacial Research’s FaceBase^61, 71, 86, 88^ Consortium (facebase.org), Record ID 3B-Y34G; DOI 10.80001/3B-Y34G. These data are additionally deposited in NCBI’s Gene Expression Omnibus^85,87^and are accessible through GEO Series accession number GSE235858. Additional data supporting the findings of this study are available from the corresponding author upon reasonable request. Images of embryos with *lacZ-*reporter activity are available from the VISTA Enhancer Browser (enhancer.lbl.gov), and raw images are available on request from the lead authors.

## Extended Data

**Extended Data Figure 1.**
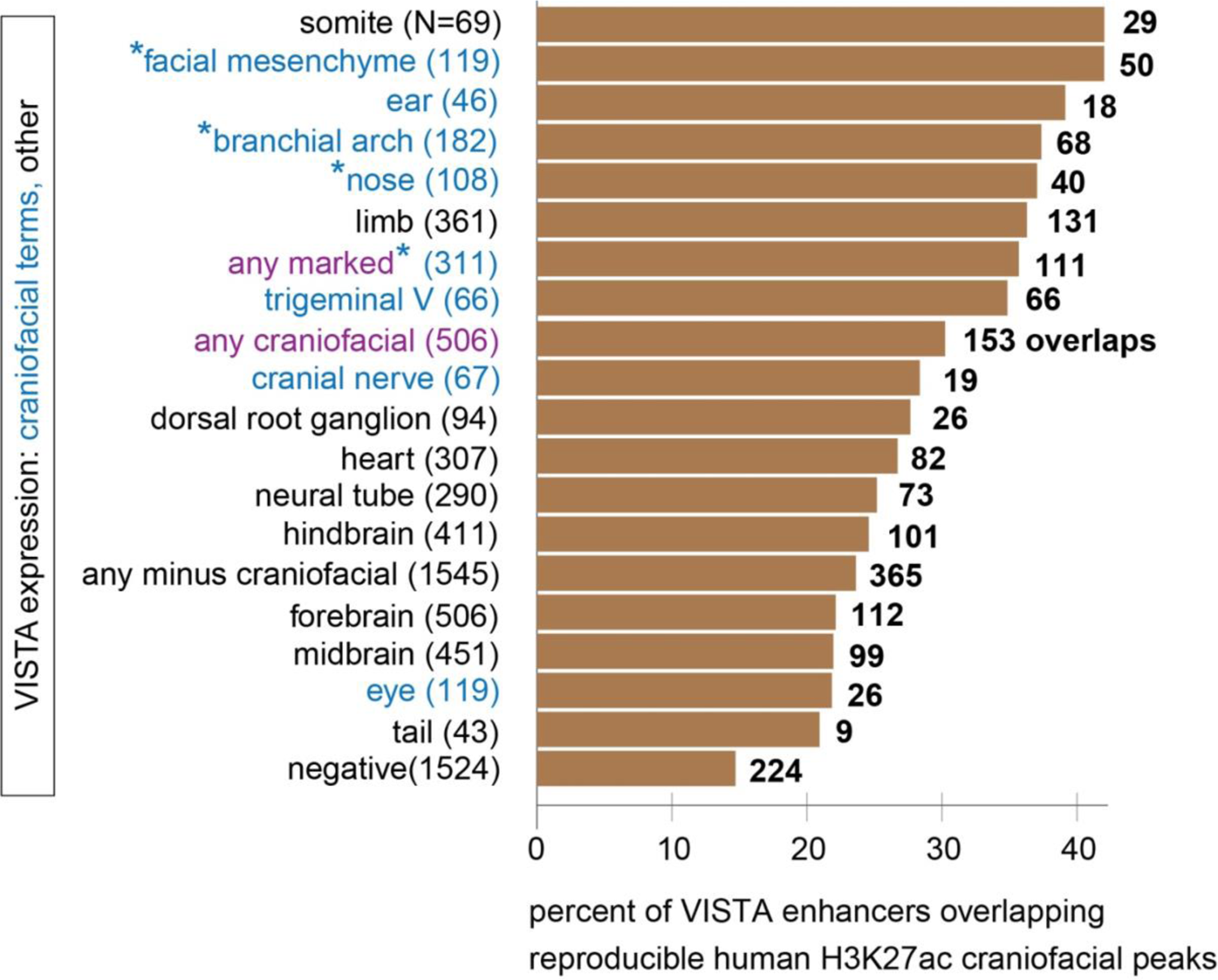
Comparison of human face enhancers and VISTA craniofacial enhancers. Y-axis shows terms for recorded expression of enhancer driven *lacZ* for elements reported in the VISTA Enhancer Browser, numbers in parentheses denote the total number (N) of such observations in the VISTA catalog. Specific craniofacial terms are denoted in blue, “any craniofacial” comprises the seven craniofacial terms shown here plus “melanocytes”. Terms with fewer than 40 elements, incl. melanocytes, are not shown individually. Bars on the right indicate the percentage of VISTA enhancers (x-axis) for the relevant expression category that overlap human developmental face enhancers in this study (n=13,983). The number in bold by each bar denotes the absolute number of such overlaps out of the total N for each expression category. See **Supplemental Table 4** for a full list of elements positive for craniofacial terms.

**Extended Data Figure 2.**
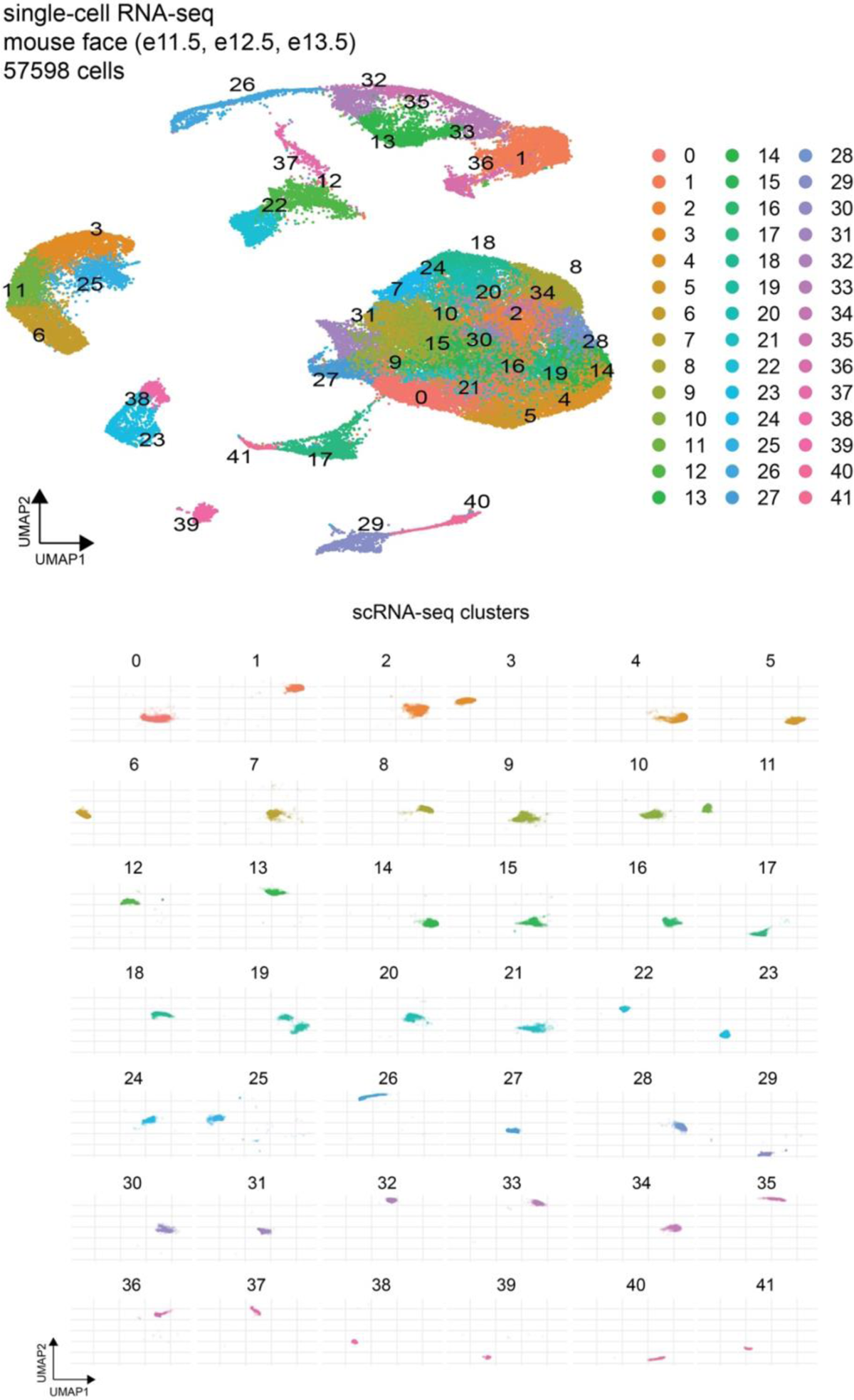
Unbiased clustering of single-cell gene expression data of the developing mouse face. For clarity, individual UMAPs are shown to demonstrate the spatial extent along x-y coordinates for each cluster (0-41) that comprises the final UMAP shown in **Figure 3**.

**Extended Data Figure 3.**
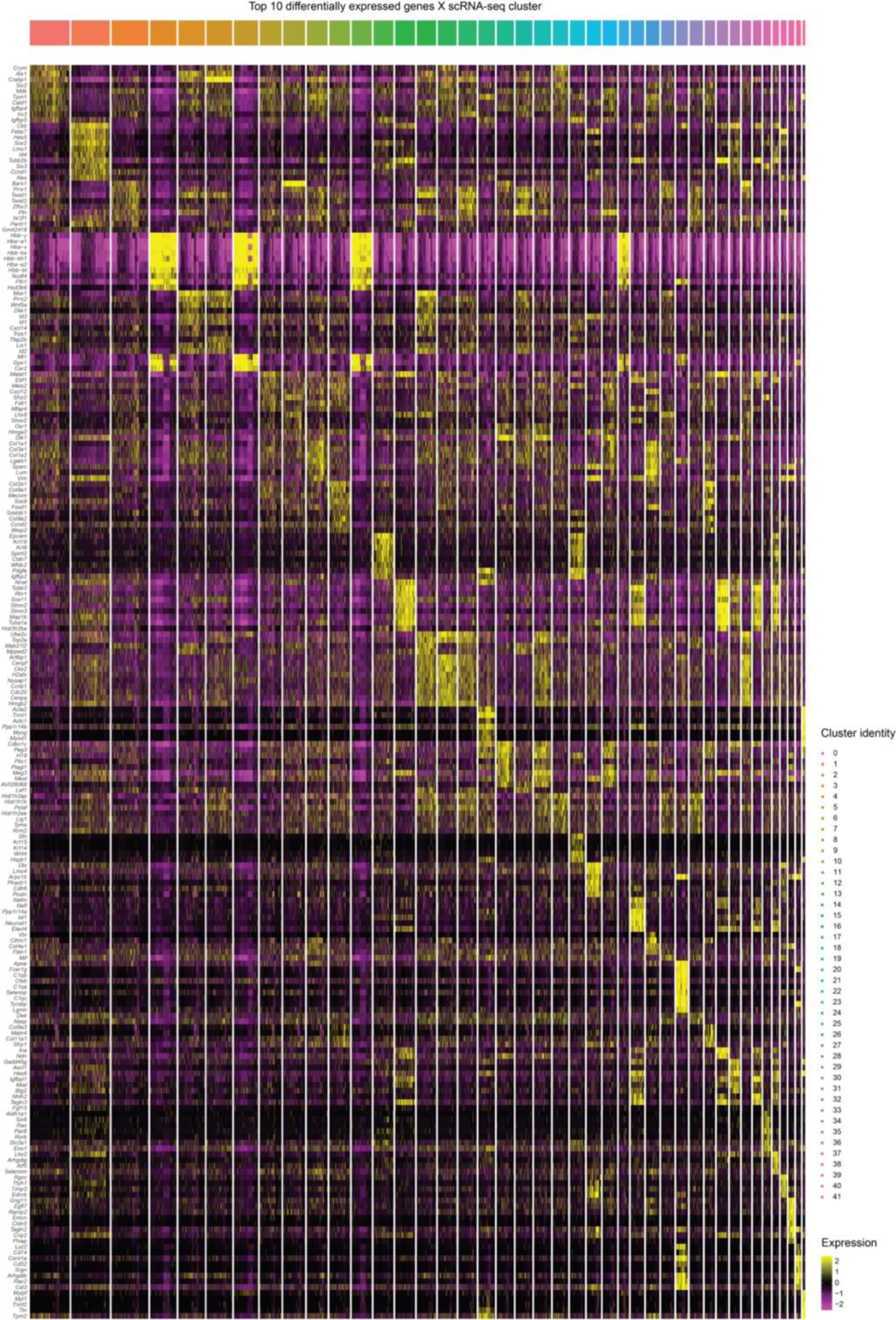
Cluster-wise marker genes in single-cell (sc) gene expression data of the developing mouse face. Heatmap shows top 10 marker genes, i.e., genes highly enriched for expression in a cluster over all clusters (y-axis) for each of the 42 original clusters (x-axis) defined in the single-cell gene expression data.

**Extended Data Figure 4.**
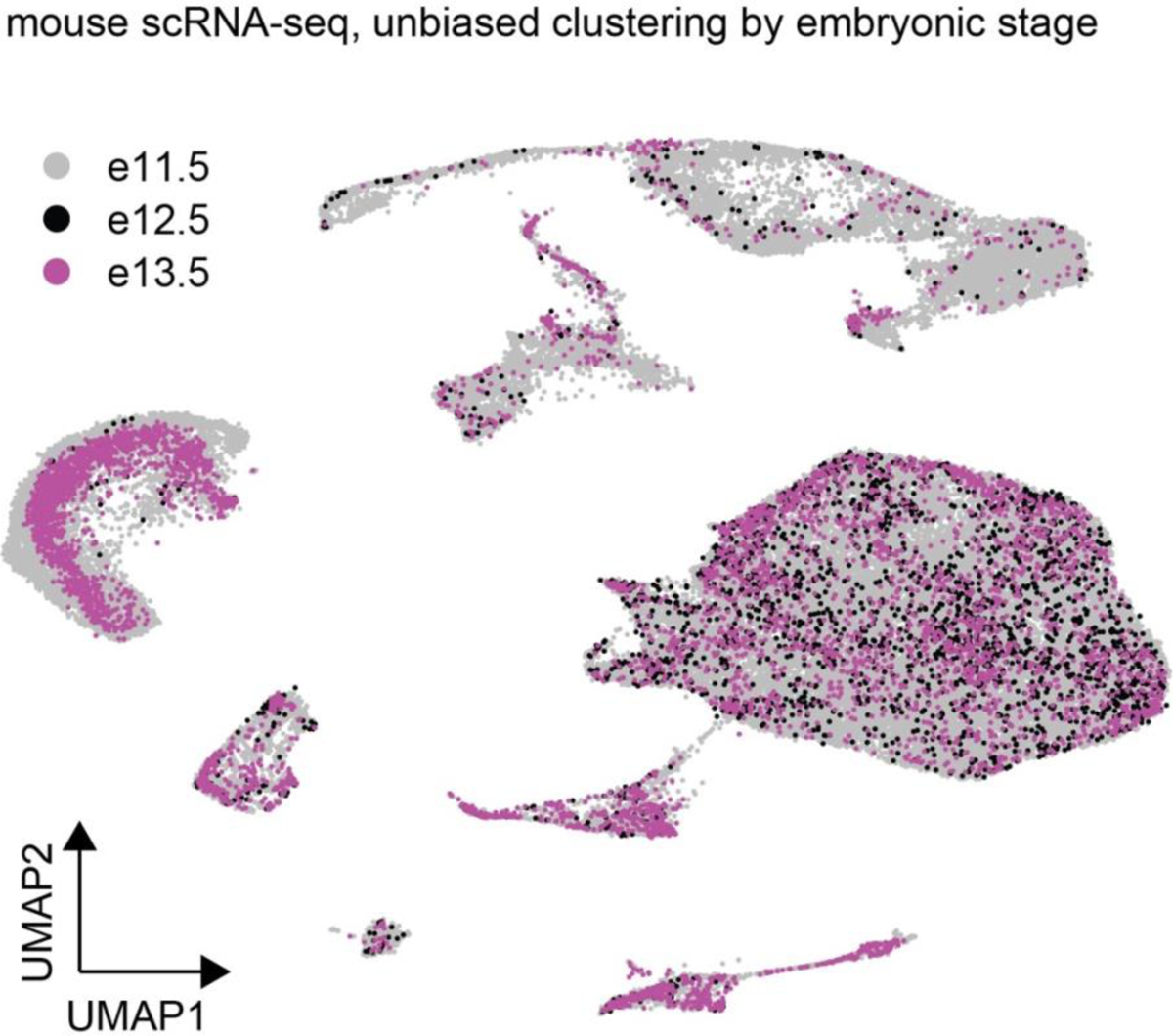
Distribution of cells in the single-cell gene expression data by mouse embryonic stage. UMAP shows distribution of cells from respective mouse embryonic stages e11.5 (gray: 49,882 cells), e12.5 (black: 2,340 cells) and e13.5 (magenta: 5,376).

**Extended Data Figure 5.**
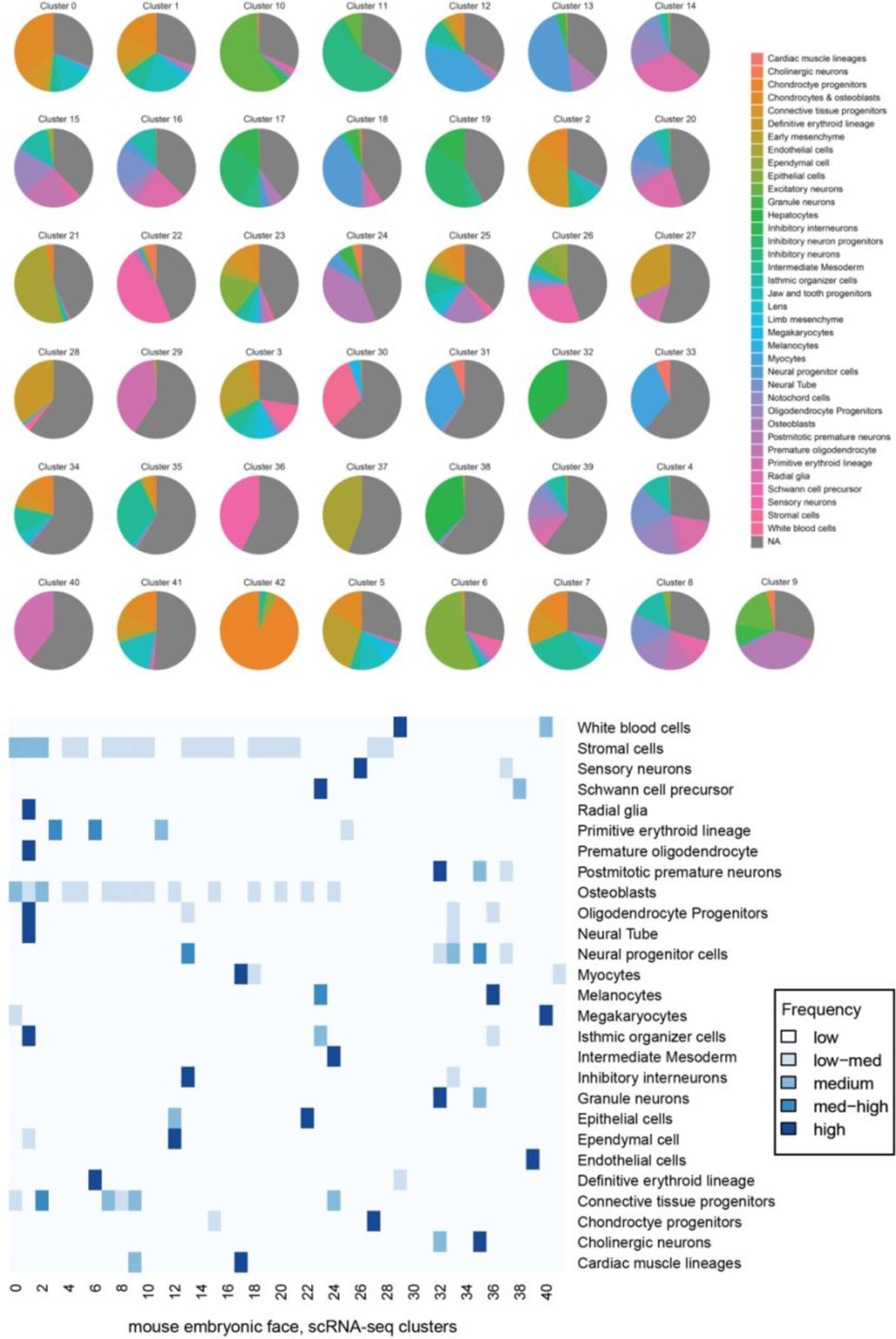
Cluster-wise proportion of cell-types in mouse single-cell gene expression data. Our single-cell gene expression data was queried to a previously published scRNA-seq large dataset of whole embryo developmental timepoints (Cao et al., 2019) using Seurat-based auto referencing as a first step for assigning cell-type identities in an unbiased manner. Heatmap shows frequency (low to high) of cell types from the reference (n=27, y-axis) that are reflected in each of the 42 clusters (x-axis). These collectively informed the first broad annotations of cell types for our *ScanFaceX* data.

**Extended Data Figure 6.**
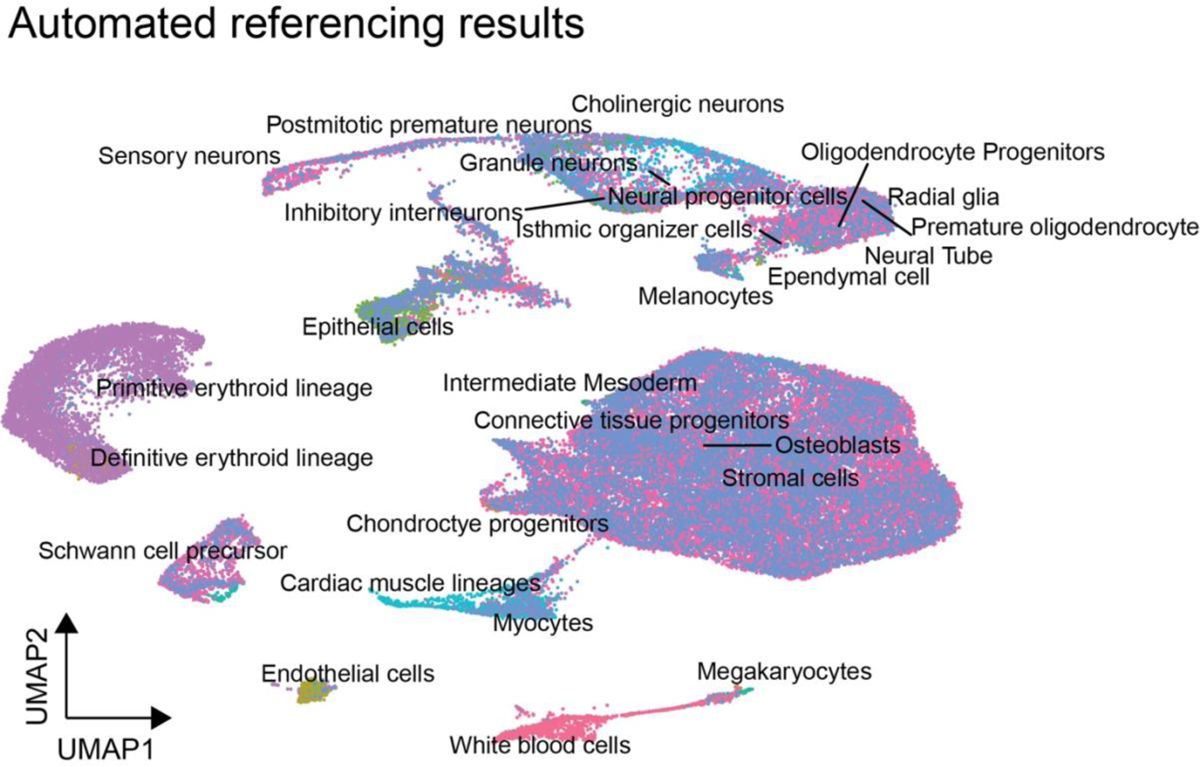
Assignment of cell-type identities in *ScanFaceX*. UMAP shows raw results from Seurat-based automated referencing and cell type annotations for *ScanFaceX* data. A down-sampled (100K cells) subset from Cao et al., 2019 whole-embryo single cell data was used as reference.

**Extended Data Figure 7.**
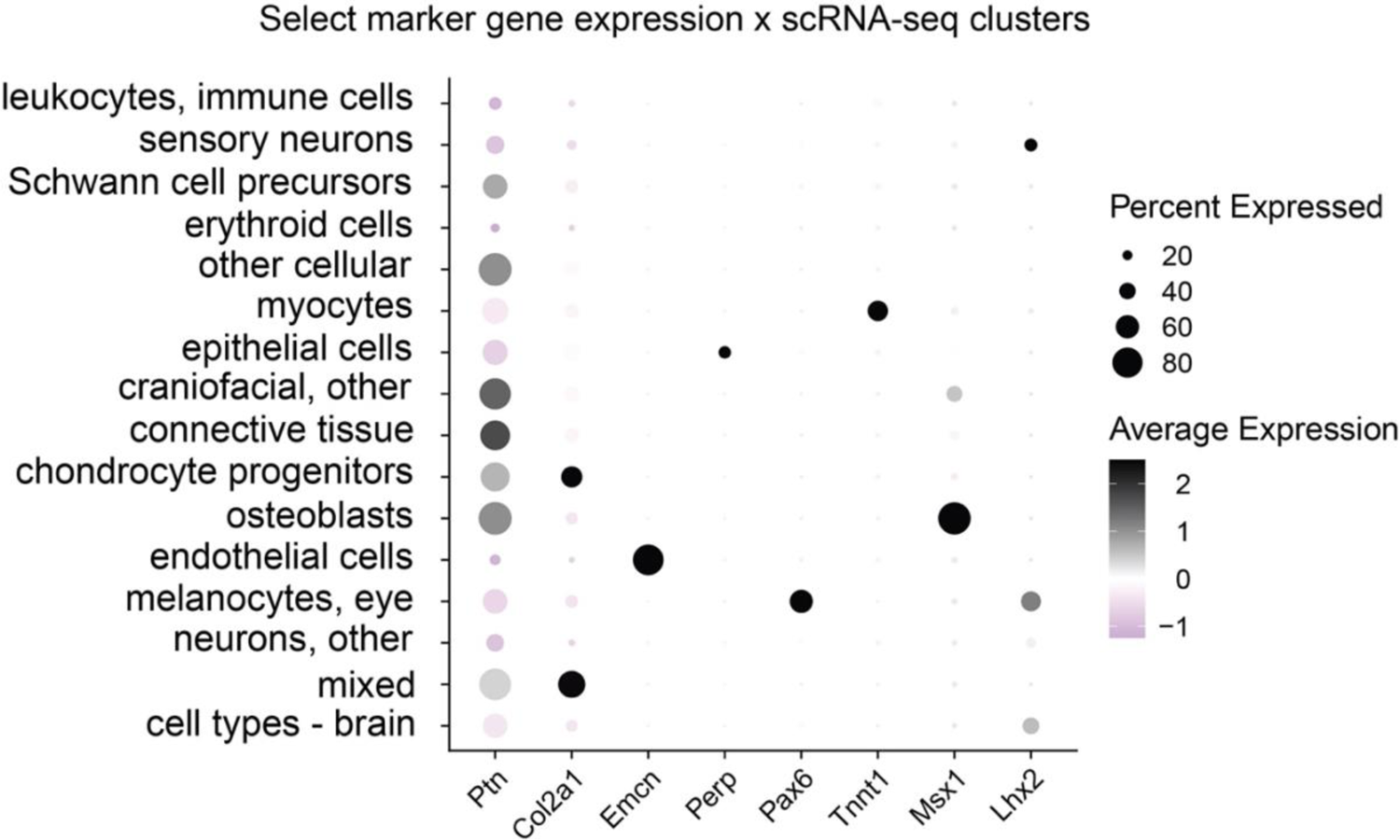
Expression of select marker genes across all clusters in *ScanFaceX*. Dot plot shows expression of select marker genes across all original clusters (consolidated per final annotations in **Supplemental Table 12**) of *ScanFaceX*, a subset of this plot is shown in main **Figure 3**. Color scale denotes low (light grey) to high (black) expression while increasing circle diameters denote corresponding higher proportion of cells within respective clusters.

**Extended Data Figure 8.**
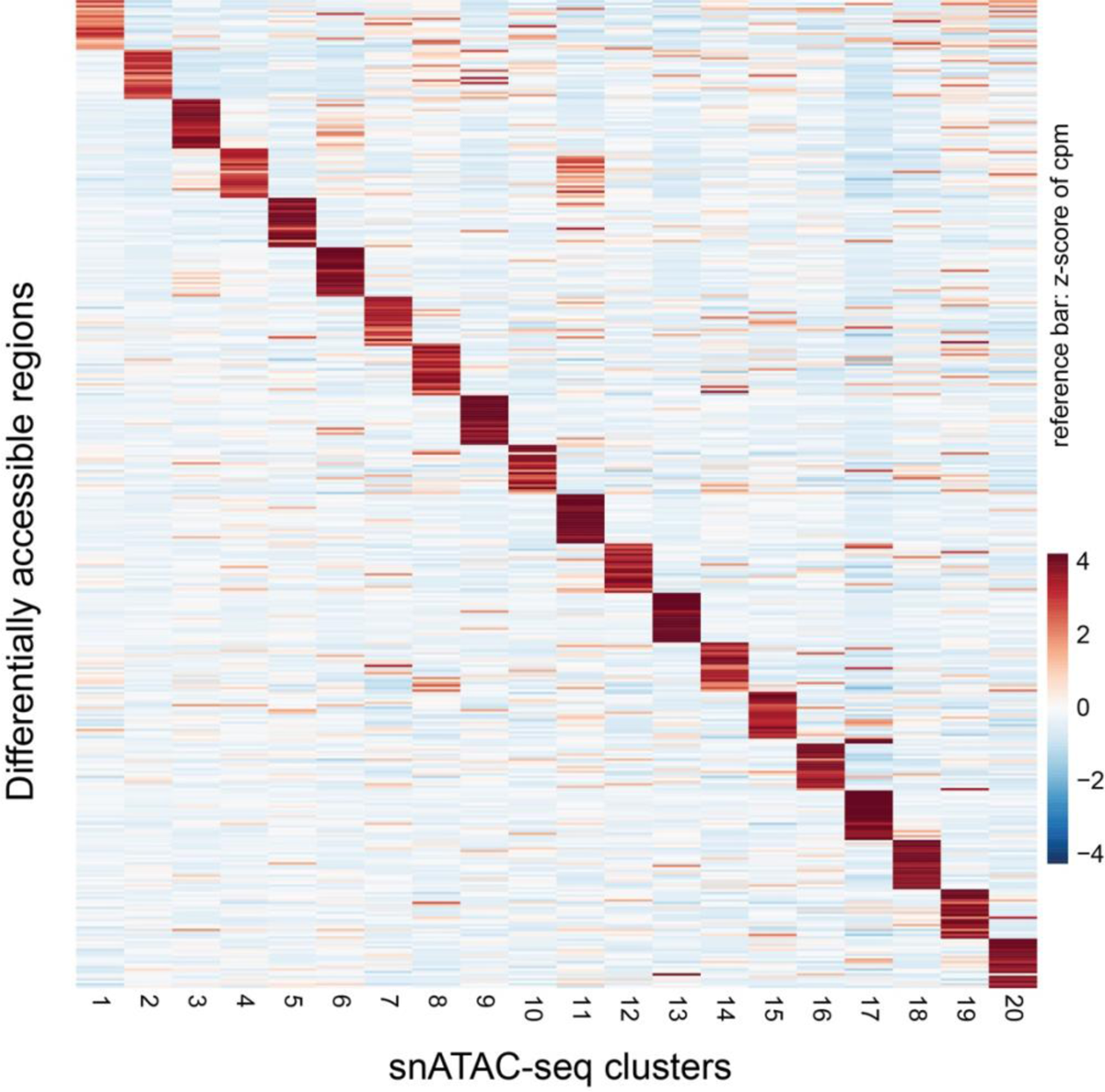
Differentially accessible regions in snATAC-seq, mouse face. Heatmap shows the top 20 DARs map exclusively to each of the 20 clusters in snATAC-seq (mouse face).

**Extended Data Figure 9.**
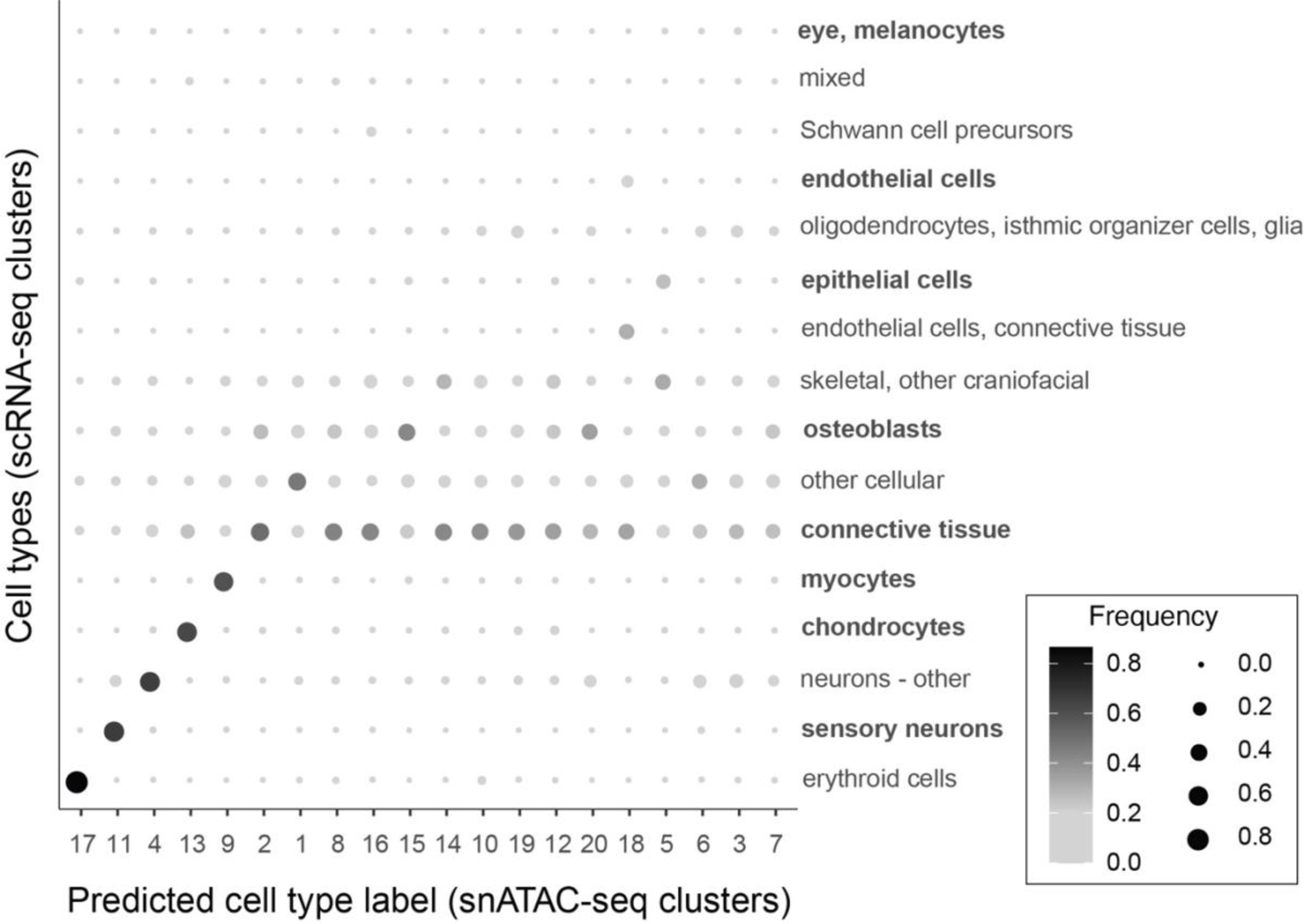
Correlation of scRNA-seq and snATAC-seq face data. Dot plot shows correlation, i.e., strength of label transfer between gene expression quantification (**scRNA-seq**; y-axis; n=16) and accessibility in TSS and gene bodies (snATAC-seq; x-axis; n=20) for integrated gene expression and open chromatin data for final annotated cell-types. Color scale denotes low (light grey) to high (black) degree of correlation while increasing circle diameters denote corresponding higher proportion of cells within correlated cell types for the respective clusters. Cell types in bold are cell types shown in **Figures 3-5**.

**Extended Figure 10.**
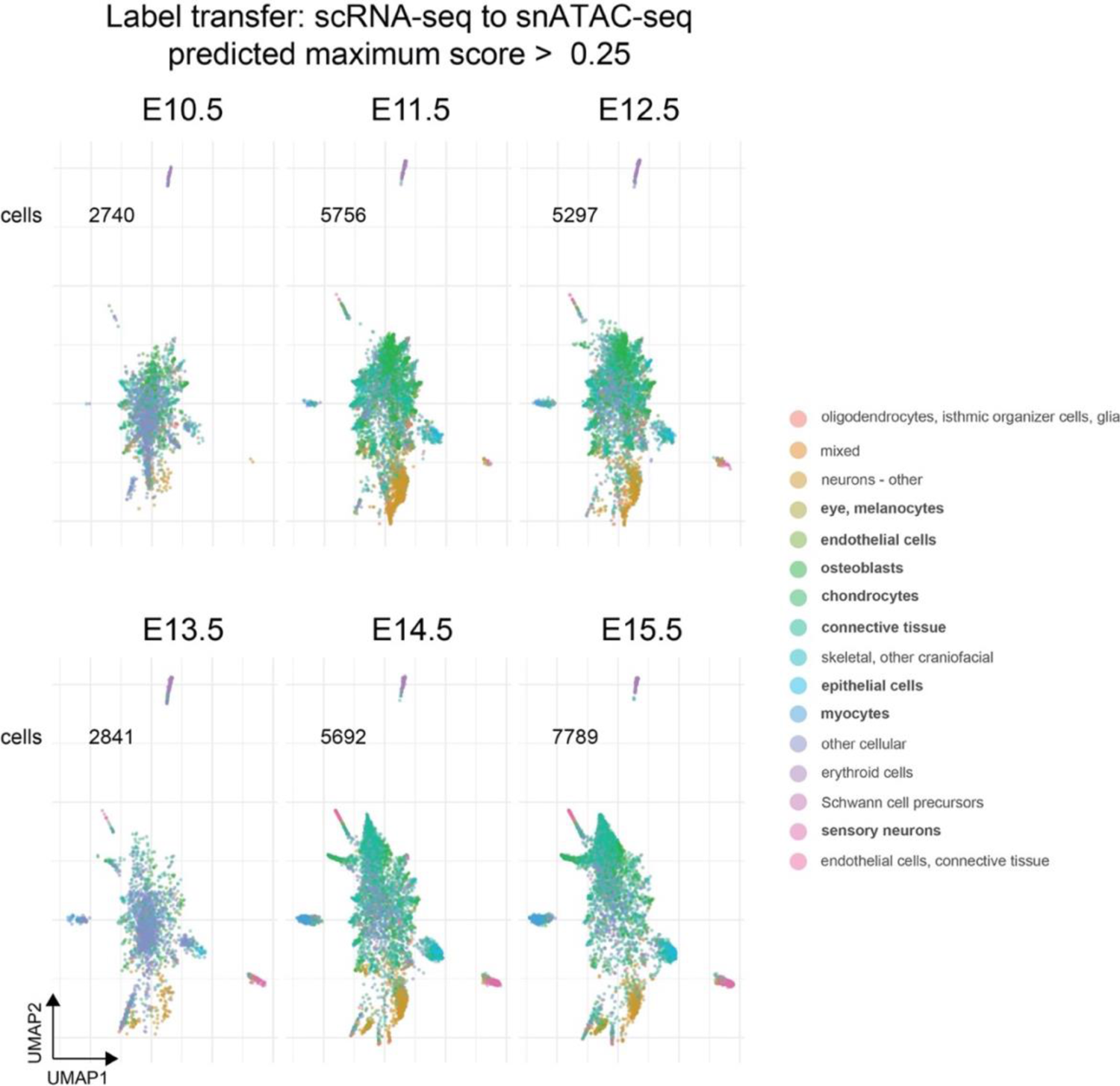
Developmental stage-wise correlation of scRNA-seq and snATAC-seq face data. Individual UMAPs show the total number of cells in our snATAC-seq assay that pass the >0.25 threshold for the predicted maximum score for label transfer between the integrated scRNA-seq and snATAC-seq datasets for 16 final cell-type annotations (key on the right).

**Extended Data Figure 11.**
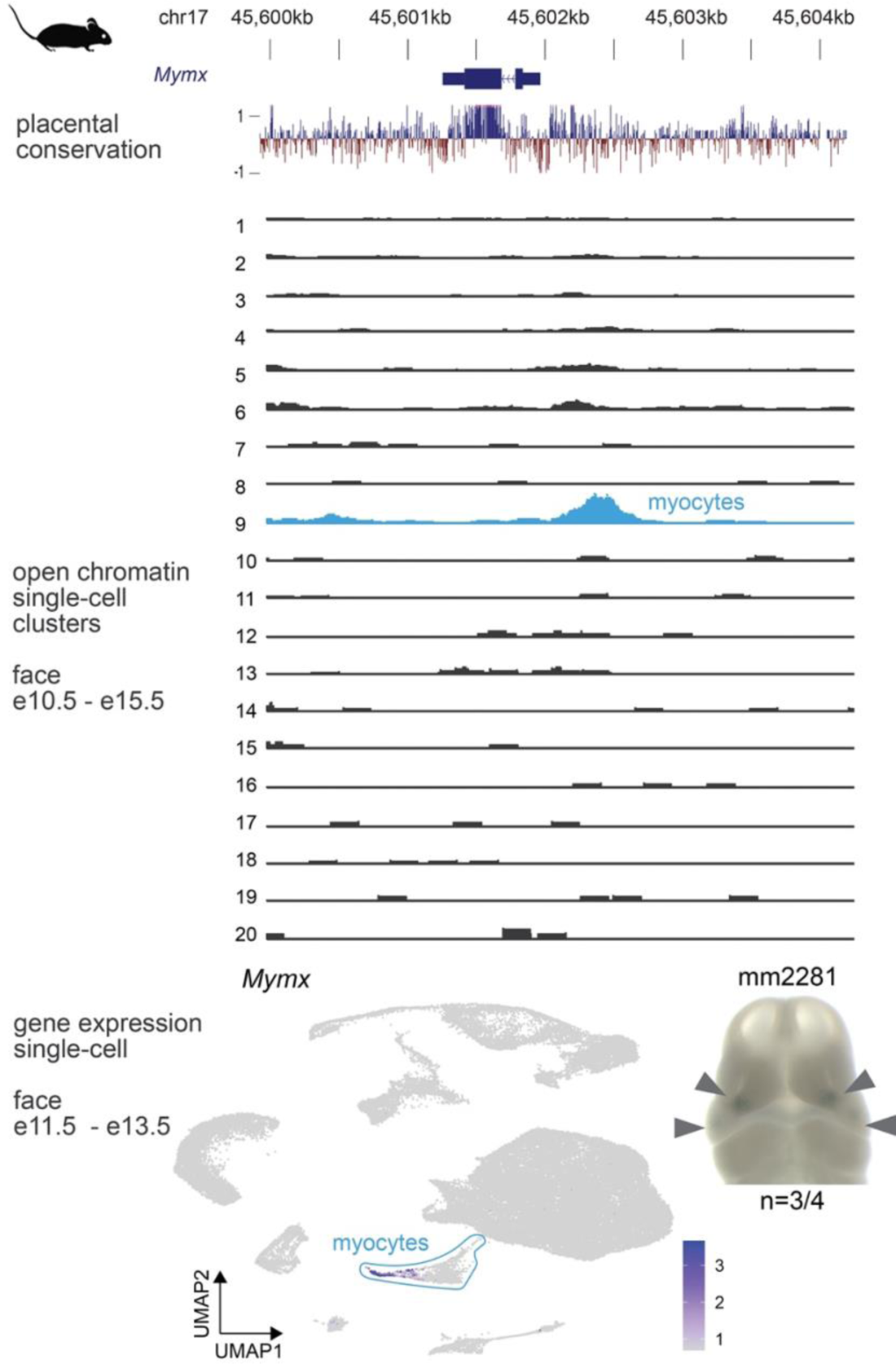
Differentially accessible regulatory regions correlate with cell-type specific signatures. The genomic context and placental conservation scores for a regulatory region near *Mymx* promoter are shown, followed by tracks for individual snATAC-seq clusters from developing mouse face tissue (e10.5 – e15.5). This region shows distinct open chromatin signature in the myocyte-specific cluster. UMAP of ScanFaceX shows expression of *Mymx* in myocytes. Image for a representative mouse embryo at e11.5 shows validated *in vivo lacZ*-reporter activity (grey arrowheads) of this enhancer. *MYMX (Myomixer)* encodes an integral membrane protein that regulates myoblast fusion, is conserved across vertebrates and *MYMX* mutations underlie an autosomal recessive disorder, Carey-Fineman-Ziter syndrome-2 (CFZS2) in humans that is characterized by weakness of the facial musculature, hypomimic facies, micrognathia, and facial dysmorphism among a range of other defects. n, reproducibility of each pattern across embryos resulting from independent transgenic integration events.

